# Protein Phosphatase PP1 Regulation of Pol II Phosphorylation is Linked to Transcription Termination and Allelic Exclusion of VSG Genes and TERRA in Trypanosomes

**DOI:** 10.1101/2023.10.21.563358

**Authors:** Rudo Kieft, Yang Zhang, Haidong Yan, Robert J. Schmitz, Robert Sabatini

## Abstract

The genomes of *Leishmania* and trypanosomes are organized into polycistronic transcription units flanked by a modified DNA base J involved in promoting RNA polymerase II (Pol II) termination. We recently characterized a *Leishmania* complex containing a J-binding protein, PP1 protein phosphatase 1, and PP1 regulatory protein (PNUTS) that controls transcription termination potentially via dephosphorylation of Pol II by PP1. While *T. brucei* contains eight PP1 isoforms, none purified with the PNUTS complex, suggesting a unique PP1-independent mechanism of termination. We now demonstrate that the PP1-binding motif of TbPNUTS is required for function in termination *in vivo* and that TbPP1-1 modulates Pol II termination in *T. brucei* involving dephosphorylation of the C-terminal domain of the large subunit of Pol II. PP1-1 knock-down results in increased cellular levels of phosphorylated large subunit of Pol II accompanied by readthrough transcription and pervasive transcription of the entire genome by Pol II, including Pol I transcribed loci that are typically silent, such as telomeric VSG expression sites involved in antigenic variation and production of TERRA RNA. These results provide important insights into the mechanism underlying Pol II transcription termination in primitive eukaryotes that rely on polycistronic transcription and maintain allelic exclusion of VSG genes.

## INTRODUCTION

Transcription can be divided into three stages: initiation, elongation, and termination. During transcription termination, RNA polymerase discontinues elongation of the RNA product and releases the DNA template. The largest subunit of the RNA Polymerase II (Pol II) complex, RPB1, contains a C-terminal domain (CTD) composed of a heptad peptide that is repeated 26 and 52 times in budding yeast and human, respectively (1). Dynamic phosphorylation of residues within this domain plays a critical role in all stages of transcription, including termination, and thus gene expression (2,3). Pol II termination of protein-coding genes in eukaryotes is tightly linked to the processing of the nascent transcript 3’-end by a mechanism referred to as “torpedo” termination. According to the “torpedo” model, cleavage of the nascent transcript at the poly(A) site provides access for the 5’-3’ RNA exonuclease Xrn2/Rat1 (human/yeast) (4–8) that eventually leads to dissociation of the polymerase from the DNA template. PP1-mediated dephosphorylation of the transcription elongation factor Spt5, resulting in deceleration of transcription downstream of poly(A) sites enhancing the torpedo dislodgment of Pol II (4,5). PP1 mediated dephosphorylation of Pol II CTD coordinates the recruitment of factors involved in 3’ end formation and termination (9). In yeast, PP1 dephosphorylation of both Pol II CTD and Spt5 is thought to orchestrate the recruitment of the termination factor Seb1 and the transition from elongation to termination and Pol II release (10).

Substrate specificity of Ser/Thr phosphatases, such as PP1, is mainly dependent on regulatory proteins, which play targeting, substrate specificity and inhibitory roles (11–13). PP1 functions in termination as part of a PTW/PP1 complex, which contains the phosphatase bound to the PP1 regulatory protein PNUTS along with Wdr82 and the DNA binding protein TOX4 (5,9,14-19). PNUTS is the scaffolding protein in the complex and mediates independent associations of PP1, Wdr82 and Tox4. PP1 binds the central regions of PNUTS via the conserved RVXF PP1-binding motif, where the second and fourth residues bury deep in two hydrophobic pockets on the PP1 surface, providing an essential stabilizing force (20). Mutation of the hydrophobic positions in the RVXF-binding motif, as it does with most PP1 interactive proteins, abolished the ability of PNUTS to bind to PP1 (20,21). Overall, the PTW/PP1 complex is a negative regulator of RNA Pol II elongation rate and plays a key role in transcription termination. Depletion of individual components in human cells, or ortholog components in yeast, leads to RNA Pol II transcription termination defects (22–26)

Very little is understood regarding the transcription cycle in trypanosomatid protozoa. In these early divergent eukaryotes, protein coding genes are arranged into polycistronic transcription units (PTUs) transcribed by Pol II (27–29). PTUs can be divergent or convergent and are separated by strand switch regions (SSRs). Pre-mRNAs are processed through trans-splicing with the addition of a 39-nucleotide spliced leader sequence to the 5’ end of mRNAs, which is coupled to the 3’ polyadenylation of the upstream transcript (30–37). This coupling of RNA processing events may prevent the generation of a 5’-P substrate for the exonuclease during 3’-end formation and thus, allow bypass of torpedo termination within the PTU. Transcription termination sites (TTSs) at the end of PTUs, for example at convergent SSRs (cSSRs), are enriched in three chromatin marks: the hypermodified DNA base J, H3V and H4.V (38–41). Base J consists of a glucosylated thymidine (42) and has been found only in the nuclear DNA of trypanosomatids, *Diplonema*, and *Euglena* (43,44). In trypanosomatids base J is found at almost all Pol II transcription termination sites (41,45). All three epigenetic marks have been shown to play a role in Pol II transcription termination (45–50). Furthermore, for several PTUs in *Trypanosoma brucei* and *Leishmania*, J/H3V are found to promote Pol II termination prior to the end of the gene cluster, leading to silencing of the downstream genes (46–48). Loss of J/H3V from these premature termination sites results in readthrough transcription and de-repression of the downstream genes.

Recently, a complex related to the human PTW/PP1 complex was identified in *Leishmania* composed of PP1-PNUTS-Wdr82 and a DNA base J-binding protein, JBP3 (referred to as the PJW/PP1 complex) (51,52). Among the eight PP1 isoforms identified in the *Leishmania* genome, only PP1-8e is found associated with the PJW/PP1 complex *in vivo*. *Leishmania* PNUTS contains a conserved RVXF PP1-binding motif (52) and alanine substitution of the hydrophobic residues in the motif disrupts PNUTS-PP1-8e association (53). Deletion of PP1-8e and JBP3 in *Leishmania* (51,53) leads to Pol II termination defects similar to the defects following the loss of base J/H3V. Purified *Leishmania* PJW/PP1 complex was able to dephosphorylate the large subunit of Pol II (53). In contrast, no Pol II dephosphosphorylation was evident using complex lacking PP1-8e. Demonstrating the PP1-8e subunit of PJW/PP1 exhibits specific phosphatase activity toward RPB1 *in vitro,* suggested that the PJW/PP1 complex is involved in PP1-mediated Pol II-CTD dephosphorylation in the *Leishmania* nucleus and linked to the readthrough transcription defects in the PP1-8e KO. Supporting a conserved PNUTS-PP1 regulatory mechanism from trypanosomatids to yeast and mammalian cells. We therefore proposed that similar to the human PTW/PP1 complex, LtPNUTS is scaffolding protein that mediates the binding of PP1, JBP3 and Wdr82, with JBP3 tethering the complex to base J-marked transcription termination sites allowing PP1-mediated dephosphorylation of Pol II and control of termination.

Presumably, a similar mechanism of Pol II termination via the PJW/PP1 complex exists in the African trypanosome, *T. brucei*. However, purification of TbPNUTS pulls down JBP3 and Wdr82 but not PP1 (52). Supporting the conserved function of the PJW complex in *T. brucei*, depletion of TbPNUTS, TbJBP3 or TbWdr82, leads to Pol II termination defects, similar to the defects following the loss of base J/H3V (52). Although the *T. brucei* genome also harbors eight PP1 isoforms and TbPNUTS contains a conserved RVxF PP1-binding motif, no obvious PP1 isoform is a homologue of the Leishmania PP1-8e (51,53) or is found associated with the PJW complex in vivo (52). The predicted LtPNUTS:PP1-8e holoenzyme complex, supported by biochemical studies, revealed that LtPNUTS binds PP1-8e using an extended RVxF-ɸ_R_-ɸɸ-Phe motif similar to that used by the human PNUTS:PP1 complex (54). These studies also indicated that PP1 isoform specificity of LtPNUTS binding involves unique sequences in PP1-8e that are required for stable LtPNUTS:PP1-8e binding. The lack of these unique PP1 sequences in all TbPP1 isoforms could therefore explain the lack of stable PP1 association during purification of the PJW/PP1 complex from *T. brucei*. However, the lack of PP1 stably associated with the PJW complex in *T. brucei* prevented the analysis of a conserved mechanism of Pol II termination in trypanosomatids and suggested the possibility of the unique PP1-independent mechanism.

Also unique for a eukaryote, *T. brucei* contains a multifunctional RNA polymerase I (Pol I) able to synthesize the typical rRNAs and a subset of abundant pre-mRNAs critical for parasite survival in the mammalian host. *T. brucei*, is transmitted by tsetse flies to the mammalian bloodstream, thereby causing human sleeping sickness. During this bloodstream form (BSF) life-stage, the exclusive expression of only one Variant Surface Glycoprotein (VSG) gene per cell and the periodic switching of the expressed VSG allows the parasite to evade the host immune system (55,56). While *T. brucei* encodes >2500 VSG genes, only one can be transcribed when located within one of ∼15 telomeric VSG expression sites (ESs). These so-called bloodstream ESs (BESs) are Pol I transcribed PTUs located adjacent to telomeres on different chromosomes. Each BES contains a VSG gene and up to 12 expression site-associated genes (ESAGS) (57,58). BESs are transcribed by Pol I that terminates within a few kilobases from the promoter except at one fully transcribed ES (59,60). This highly stringent monoalleleic exclusion ensures that only one ES is transcribed at a time (61–63). In addition, Pol I readthrough transcription of the single active ES into the downstream telomeric repeats in BSF trypanosomes result in the production of a telomere repeat-containing long, non-coding RNA called TERRA (64,65). Once the parasites are back in the fly and migrate to the salivary gland they differentiate into metacyclic forms. To preadapt for infection of the mammalian host, metacyclic forms are also covered by a VSG coat that is Pol I transcribed from one of several specific telomeric metacyclic expression sites (MES) (66,67). Interestingly, base J and H3V are enriched in the silent ESs (38,41) and loss of H3V/J (47,49,57) as well as depletion of components of the PJW complex (52) lead to de-repression of silent VSG ESs.

We now demonstrate that the PP1-binding motif of TbPNUTS is required for function in Pol II transcription termination *in vivo*, suggesting that a PP1-PNUTS complex is involved in Pol II transcription in *T. brucei*. Using RNAi to screen all the functional PP1 isoforms, we show that the specific depletion of TbPP1-1 leads to readthrough transcription and pervasive transcription of the entire genome by Pol II, including silent Pol I transcribed loci, such as the telomeric VSG expression sites. Therefore, the readthrough transcription defect results in the loss of mono-allelic exclusion of VSG expression and increased levels of TERRA. PP1-1 knock-down also results in increased cellular levels of phosphorylated large subunit of Pol II. We propose a model whereby TbPP1-1 activity, as a component of the PJW/PP1 complex, controls Pol II CTD phosphorylation status and termination of transcription of PTUs in the chromosome core of *T. brucei*, preventing pervasive transcription and expression of the silent sub-telomeric VSG arrays and Pol I telomeric VSG ESs. These results provide important insights into the mechanism underlying transcription termination in primitive eukaryotes that rely on polycistronic transcription and the need to bypass termination signals at the end of individual genes and suggests a role of Pol II regulation in the control of VSG gene allelic exclusion.

## MATERIALS AND METHODS

### Parasite culture

Bloodstream form *T. b. brucei* “single marker cells (SMC)”, expressing T7 RNA polymerase and the Tet repressor (68) were used in these studies and cultured in HMI-11 medium, supplemented with 10 % Tetracycline free FBS at 37°C. Transfections were performed using the Amaxa electroporation system (Human T Cell Nucleofactor Kit, program X-001) and clonal cell lines obtained as described (52). Where appropriate, the following drug concentrations were used: 2 μg/ml G418, 2.5 μg/ml Hygromycin, 2.5 μg/ml Phleomycin, 2 μg/ml Blasticidin. Procyclic form *T. b. brucei* 29-13 were cultured in SDM79 medium at 27°C.

### DNA constructs and cell line generation

Tagging the C-terminus of the PP1 with a triple HA or Myc epitope was performed using a PCR based approach with the pMOTag4H / pMOTag3M construct as previously described (52). For Tetracycline regulated expression of RNAi resistant PNUTS in *T. brucei*, DNA was synthesized by GeneArt to recode the entire ORF (ensuring there is at least one silent mutation for every 12 nucleotides) and include a C-terminal Protein A tag. The synthesized DNA was sub-cloned into the HindIII and BamHI sites of the pLew100V5 plasmid by Gibson cloning. The final construct (WT PNUTS-ProtA-pLew100V5), was linearized with NotI prior to transfection. A subsequent clone was made by swapping out the Protein A tag with a FLAG tag. The final construct (WT PNUTS-FLAG-pLew100V5) was linearized with NotI prior to transfection. The desired PNUTS mutant in the RVXF motif (converting the RISW motif to RASA) was generated by oligonucleotide-mediated site-directed mutagenesis (QuikChange II XL Site-Directed Mutagenesis Kit, Agilent Technologies) following the manufacturer’s instructions. To allow induction of RNAi resistant untagged version of PP1-1, PP1-6 and PP1-7, DNA for each was recoded and cloned into the pLew100V5 plasmid and transfected as described for PNUTS above. To induce PNUTS or PP1 expression, Tetracycline was added at 2 ug/ml. All final constructs were sequenced prior to electroporation.

### RNAi analysis

Conditional silencing of PNUTS was performed as previously described (52). For conditional PP1 silencing experiments, a clade specific fragment (69) of the ORF was integrated into the BamHI site of the p2T7-177 vector. I-SceI linearized p2T7-177 constructs were transfected into BF SMC for targeted integration into the 177 bp repeat locus. RNAi resistant PP1 rescue constructs were generated and transfected as described above. All final constructs were sequenced prior to transfection. RNAi and/or expression of the recoded PP1 was induced with 2 μg/ml Tetracycline and growth was monitored daily in triplicate.

### Transcription inhibition

BF cells were incubated with 1 μM BMH-21 for 15 min or 5 μg/ml Actinomycin D for 2 hrs at 37°C in medium. PC cells were incubated with 1 μM BMH-21 for 15 or 120 min, or 5 μg/ml Actinomycin D for 2 hrs at 27°C in medium. Control incubations were performed using DMSO at similar concentrations. Post incubations, cells were harvested and immediately lysed in Tripure Isolation Reagent (Roche) and stored at −80°C until processing.

### Northern analysis

Total RNA was isolated with Tripure Isolation Reagent. For Northern blot analysis, 5 μg of Turbo DNase treated total RNA was separated on a 1.2 % agarose gel (1 x MOPS, 2.8 % formaldehyde), transferred to a nitro cellulose filter in 20 X SSC and UV cross-linked. Radiolabeled oligonucleotides were generated and hybridized in an aqueous hybridization buffer with 100 μg/ml yeast tRNA at 60°C. A random primer labeled β-Tubulin probe was generated and hybridized in a 40 % (v/v) formamide hybridization mix with the addition of 10% (w/v) dextran sulfate and 100 μg/ml yeast tRNA at 42°C. Final washes were performed with 0.3 X SSC / 0.1 % SDS at 60°C for 30 min.

For RNA dot blot analysis, 2 μg of Turbo DNase treated total RNA was incubated with or without 25 U RNase One (Promega) and 25 μg RNaseA (Thermo Scientific) for 1 hr at 37°C. An equal volume of 2 x loading buffer (50 % formamide, 5.6 % formaldehyde, 2 x MOPS) was added, incubated at 80°C for 10 min, snap-cooled on ice, applied to a dot blot apparatus and UV cross-linked.

### RT-PCR analysis

Total RNA was isolated with Tripure Isolation Reagent. cDNA was generated from 0.5 - 2 μg Turbo^TM^ DNase (ThermoFisher) treated total RNA with Superscript^TM^ III (ThermoFisher) according to the manufacturer’s instructions with either random hexamers, oligo dT primers or strand specific oligonucleotides. Strand specific RT reactions were performed with the strand specific oligonucleotide and an antisense Asf I oligonucleotide. For regular RT-PCR, equal amounts of (strand specific) cDNA was used with Ready Go Taq Polymerase (Promega). To ensure specific DNA was amplified, a minus RT control reaction was used.

For the RT-PCR analysis to determine the origin of TERRA, RNA isolation was performed the same way as described above, except DNase I digestions were repeated two additional times. Reverse transcription was performed using TELC20 or a random hexamer as the primer and PCR amplified as previously described (65). To reduce endogenous priming by the telomeric primer TELC20 in the RT reaction, the denaturation step was performed by heating to 90°C for 1 min followed by cooling to 55°C at 0.5 °C / sec. For PCR amplification using primers specific a sequence at Chromosome 11 subtelomere the number of cycles were increased from 25 to 28.

### Quantitative RT-PCR analysis

Total RNA was isolated and Turbo^TM^ DNase treated as described above. Quantification of Superscript^TM^ III generated cDNA was performed using an iCycler with an iQ5 multicolor real-time PCR detection system (Bio-Rad). Triplicate cDNA’s were analyzed and normalized to Asf I cDNA. qPCR oligonucleotide primers combos were designed using Integrated DNA Technologies software. cDNA reactions were diluted 10-fold and 5 µl was analyzed. A 15 µl reaction mixture contained 4.5 pmol sense and antisense primer, 7.5 µl 2X iQ SYBR green super mix (Bio-Rad Laboratories). Standard curves were prepared for each gene using 5-fold dilutions of a known quantity (100 ng/µl) of WT gDNA. The quantities were calculated using iQ5 optical detection system software.

### Western analysis

For Western analysis, 5 x 10^6^ – 1 x 10^7^ cell equivalents were analyzed on PAA / SDS gels and probed with anti-HA antibodies (Sigma, rat 3F10, 1:1000), anti-Myc antibodies (ThermoFisher, mouse 9E10, 1:2000), anti-eF1α antibodies (Sigma, mouse CBP-KK1, 1:20.000), anti-Protein A antibodies (Sigma, P3775, 1:5000). IRDye secondary antibodies (800CW and 680RD) were used (1:10.000). Antibody incubations were performed using Intercept Blocking Buffer; gels were scanned with an Odyssey imager and analyzed with Image Studio software (LI-COR).

### Strand-specific RNA-seq library construction

For mRNA-seq, total RNA was isolated from *T. brucei* RNAi cultures grown in presence or absence of Tetracycline for two days using TriPure. Six mRNA-seq libraries were constructed for each of the PP1 RNAi cell lines (triplicate samples for plus and minus Tetracycline) using Illumina TruSeq Stranded RNA LT Kit following the manufacturer’s instructions with limited modifications. The starting quantity of total RNA was adjusted to 1.3 µg, and all volumes were reduced to a third of the described quantity. High throughput sequencing was performed on an Illumina NovaSeq 6000 instrument.

### RNA-seq analysis

Raw reads from mRNA-seq were first trimmed using Trim Galore! (https://www.bioinformatics.babraham.ac.uk/projects/trim_galore/) with default settings (v0.6.5). Remaining reads were locally aligned to the *T. brucei* Lister v427 genome assembly downloaded from TriTrypDB (v59; (70)) and the Lister 427 BES sequences (58) using Bowtie2 version 2.4.5 (71). With non-default settings (sensitive local) and further processed with SAMtools version 1.16.1 (72). For each sample, FeatureCounts (v2.0.1; (73)) was used to count reads for each reference transcript annotation, followed by normalization/variance stabilization using DESeq2 (v1.38.3; (74)). Only sense reads for a given transcript with at least one read mapping across all the libraries were considered. Differential gene expression was conducted using DESeq2 by comparing PP1 RNAi samples with and without tetracycline in triplicate (log2 fold change and differential expression test statistics can be found in Table S1 and S2).

To compare Tetracycline-treatment fold changes for specific strands genome-wide, we counted reads from each strand in 5000bp bins with a 1000bp step. The log2 fold changes between Tet-untreated and Tet-treated RNAi data were plotted for chromosome 9 (as a representative) and all the BESs.

To analyze transcription defects at 3’- and 5’-end of PTUs, reads mapping to TTS (cSSRs) were counted and reads per kilobase per million mapped reads (RPKM) values were generated. Similar to what we have previously done (46–48), lists of cSSRs were generated computationally as defined regions where coding strands switch based on the transcriptome. Several SSRs located at subtelomeres were not included due to ambigious nature of gene organization. SSRs and the 5-kb flanking regions were analyzed with DeepTools (v3.2.1; (75)) using 100bp bins flanking SSRs and dividing each SSR into 50 equally sized bins.

## RESULTS

### PP1-1 functions in RNA Pol II transcription termination

There are eight PP1 homologs in the *T. brucei* genome (Figure S1A). Proteomic analysis indicates that seven of these homologs, all except the PP1-8 isotype, are expressed to varying degrees in both PC and BS form *T. brucei* (Figure S1B) (76). The PP1-8 gene is truncated at the N- and C-termini of the catalytic subunit and lacks the sixth manganese-coordinating residue in the active site (Figure S2) and thus, even if expressed would result in a non-functional enzyme. Consistent with the lack of PP1-8 expression in WT trypanosomes, HA-tagging the endogenous PP1-8 locus failed to detect PP1-8-HA protein by western blot (unpublished data). To explore the role of PP1 in transcriptional control we utilized RNAi to knockdown (KD) the expression of all seven functional PP1 isotypes in bloodstream form of *T. brucei*. Due to the high sequence identities between PP1-2 and PP1-3 (95%), between PP1-4 and PP1-5 (99%) and between PP1-5 and PP1-6 (95%), it was possible to use one partial DNA sequence from PP1-2 to KD both PP1-2 and PP1-3 and another sequence to KD PP1-4, PP1-5, and PP1-6 simultaneously. Using this approach, it was previously shown that RNAi induction significantly diminished corresponding PP1 mRNA levels (69). A specific partial sequence from PP1-1 and other from PP1-7 was designed to KD the corresponding PP1 isotype. The effects of RNAi on expression of the seven PP1s were examined by western blot using endogenously tagged PP1. Induction of RNAi against PP1 isotypes 1 and 4-6, and KD of protein levels from 50-80% (Figure 1A) leads to reduced parasite growth (Figure S3A), indicating that the proteins are important for normal cell proliferation in BSF *T. brucei*. The KD of PP1 isotypes 2 and 3, and corresponding 25% reduction in protein level, did not result in a significant growth defect. The KD appears to be specific, since ablation of PP1-1 had little to no effect on levels of PP1-6 and PP1-2 protein (Figure S3B). Overall, the results indicate that, after initiating the RNAi for 2 days, levels of the mRNAs and protein aimed by the RNAi diminished significantly.

**Figure 1.**
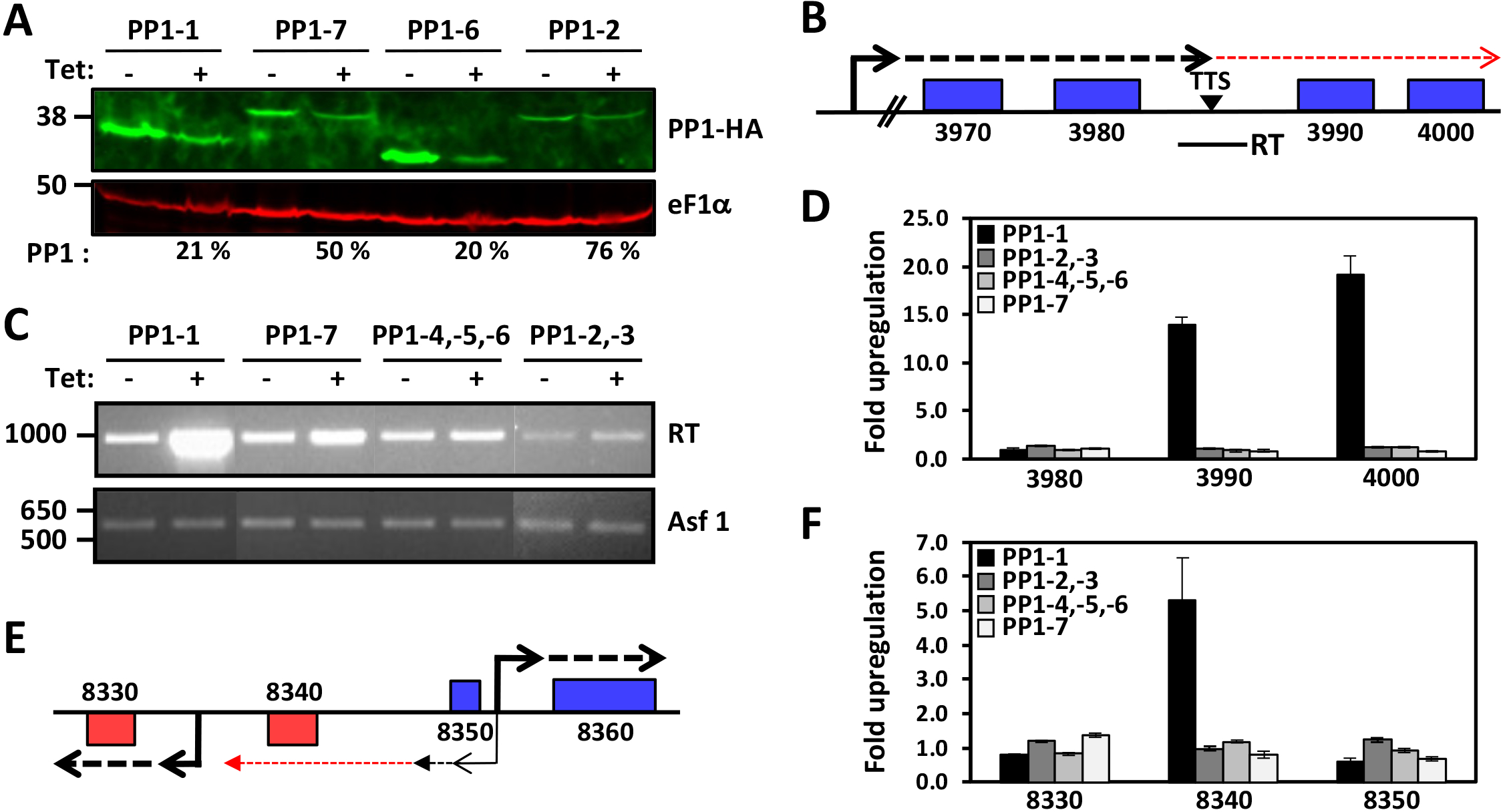
PP1-1 isoform is involved in Pol II transcription termination in *T. brucei*. (A) Depletion of PP1 isoforms upon Tet induction of RNAi for 24 hrs by anti-HA western blot analysis. Quantitation of PP1 KD, percent remaining after induction of RNAi, is indicated below. The PP1 isotype indicated represents the PP1 that is endogenously tagged and targeted by the RNAi construct. Thus, while the PP1-4,-5,-6 RNAi construct targets all three homologous RNAs, only PP1-6 protein levels are analyzed here. Probing the same blot for eF1α provides a loading control. (B-D) Analysis of Pol II termination. (B) Schematic representation of a termination site on chromosome 5 where Pol II has been shown to terminate prior to the last two genes (Tb927.5.3990 and Tb927.5.4000) in the PTU. The dashed arrow indicates readthrough transcription past the termination site (TTS) that is regulated by base J, H3V and the PJW complex. RT; Solid line below indicates the region analyzed by RT-PCR to detect readthrough (RT) RNA. (C) RT-PCR analysis of nascent RNA from the indicated clade specific PP1 depletion. cDNA was synthesized after 24 hr induction of RNAi using random hexamers and PCR was performed using the primers to amplify RT indicated in (B). Asf1 represents a PTU internal Pol II transcribed single copy gene as a loading control. (D) RT-qPCR analysis of mRNA transcript changes in the indicated PP1 isoform RNAi. RT-qPCR analysis was performed for the genes numbered according to the ORF map. Transcripts were normalized against Asf1 and fold changes (- and + Tetracycline) are plotted as the average and standard deviation of two biological replicates analyzed in triplicate. (E and F) TbPP1-1 affects early termination of antisense transcription at TSSs. (E) Representative region of chromosome 10 illustrating bi-directional transcription at a TSS. Thick black arrows indicate the start site and direction of sense transcription of the divergent PTUs. Thin black arrow indicates bi-directional non-productive transcription that typically terminates early. Red arrow indicates termination defect of the ‘antisense transcription’ upon depletion of PNUTS, Wdr82 or JBP3, resulting in de-repression of the annotated 8340 gene on the bottom strand (52). (F) mRNA transcript changes in the indicated PP1 isoform RNAi by RT-qPCR. RT-qPCR analysis was performed for the genes numbered according to the ORF map. Transcripts were normalized against Asf1 and fold changes are plotted as the average and standard deviation of two biological replicates analyzed in triplicate.

To examine the role of the PP1 isotypes in Pol II transcription termination in *T. brucei* we first used an RT-PCR assay. We have previously shown that base J and H3V are present at termination sites within a PTU where loss of either epigenetic mark results in readthrough transcription and increased expression of downstream genes (46–48). These PTU internal termination sites provide a convenient and sensitive analysis of Pol II transcription termination. For example, base J and H3V epigenetic marks and components of the PJW complex (PNUTS, Wdr82 and JBP3) are involved in terminating transcription upstream of the last two genes (VSG; Tb927.5.3990 and Hypothetical protein; Tb927.5.4000) in a PTU on chromosome 5 (Figure 1B) where defects in the termination mechanism leads to readthrough transcription and detection of nascent RNA that extends beyond the termination site (46,47,52). We have previously shown that an RNA species extending beyond this termination site following the loss of base J is indicative of readthrough transcription due to continued Pol II transcription elongation through the termination zone (46). The presence of an open reading frame downstream of the termination site allows an additional measure of readthrough where nascent RNA is processed to stable capped and polyadenylated mRNA species. As such, the loss of either epigenetic mark, or PJW complex component, in *T. brucei* leads to generation of nascent RNA extending beyond the termination site and expression of the two downstream genes (46,47,52). We now show that the specific ablation of PP1-1 leads to readthrough transcription at the representative PTU internal termination site on chromosome 5. RT-PCR using oligos immediately downstream of the termination site (see diagram, Figure 1B) detects increased RNA levels upon ablation of PP1-1 (Figure 1C). Consistent with readthrough transcription, both genes downstream of the termination site are significantly de-repressed upon the ablation of PP1-1, in contrast to genes upstream (Figure 1D). In contrast, no significant termination defects are detected upon ablation of the other PP1 isotypes (Figure 1 C and D). To analyze another termination site, we assayed for the accumulation of transcripts upstream of a transcription start site (TSS). We have previously shown that the PJW complex also terminates ‘antisense’ transcription from bi-directional initiation sites in *T. brucei (52)*. In some cases, defects in termination at these sites leads to expression of genes present upstream of the start site that are silent in WT cells. A specific example, shown in Figure 1E, includes a region on chromosome 10 where a gene (Tb927.10.8340) located between the two divergent TSSs is specifically upregulated in the PJW mutant (52). RT-PCR analysis also shows the specific upregulation of this gene ∼5-fold following the loss of PP1-1 versus genes in the adjacent PTU (Figure 1F). In contrast, ablation of the other isotypes had no significant effect. Not only do these results describe a specific function for the PP1-1 isoform in transcription, but the essential nature of the other isoforms provides a convenient negative control. Despite the significant reduction in parasite growth observed after knockdown of the essential PP1-4 and PP1-5 and PP1-6 isotypes (Figure S3A), there was little significant increase in readthrough RNAs (Figure 1B-F). This argues that the termination defects observed after the induction of TbPP1-1 RNAi is indeed a specific phenotype related to PP1-1 rather than a secondary effect of dying cells. Re-expression of an RNAi resistant PP1-1 mRNA is able to fully rescue the PP1-1 KD associated termination defect (Figure S4) and growth defect (Figure S5). This helps ensure that the phenotype was specific to the PP1-1 mutation and not due to off target effects. None of the other RNAi mutants have a defect in readthrough transcription, suggesting that none of the PP1 isotypes other than PP1-1 is involved in controlling Pol II transcription termination. However, since the KD were not total, there remains the possibility that the residual presence of PP1 could be sufficient to allow normal Pol II transcription. Interestingly, over-expression of PP1-6 is able to partially rescue the PP1-1 mutant termination defect (Figure S4). The ability of PP1-7 to rescue the PP1-1 KD is difficult to determine since over-expression of PP1-7 in WT cells alone leads to significant levels of readthrough transcription (Figure S4).

To further explore the role of the PP1-1 in transcription termination and mRNA gene expression genome wide, we performed stranded mRNA-seq to compare the expression profiles of PP1-1 RNAi cells with and without Tetracycline induction. Since PP1-1 and PP1-7 represent isotypes that have been localized to the nucleus (77), we included the PP1-7 RNAi as a control. First, we analyzed the read coverage for 5kb on either side of the termination sites at convergent strand-switch regions (cSSRs) where the 3’ termini of two PTUs converge in the *T. brucei* genome. As expected, in WT cells the mean-normalized coverage on the top strand decreased sharply at the transcription termination site (TTS) of cSSRs (Figure 2A). However, when PP1-1 is ablated, the read coverage downstream of the TTS was significantly higher (*p*-value=2.900e-8), suggesting that loss of PP1-1 resulted in significant transcriptional readthrough. Analysis of the bottom strand data reveals similar significant differences between WT and the PP1-1 mutant, where readthrough from the convergent PTU results in increased antisense transcripts past the TTS into the adjacent PTU (Figure S6). Quantitation of readthrough transcription at cSSRs genome-wide, by measuring the change in total sense and antisense reads within the cSSR, indicates a 3-fold increase upon KD of PP1-1 (*p*-value=1.118e-6) (Figure S7). In contrast, no readthrough defects are detected in the PP1-7 KD by measuring the read coverage downstream of the TTS (*p*-value=0.621) (Figures 2A and S6B) or by measuring the change in total reads within the cSSR (*p*-value=0.9128) (Figure S7). Figure 2B-D shows coverage profiles corresponding to three representative termination sites. At a cSSR on chromosome 10 (cSSR 10.4) we see that the KD of PP1-1 leads to readthrough transcription on the top strand and, to a lesser extent, the bottom strand into the adjacent PTU (Figure 2B). A Pol I transcribed EP1 locus embedded at cSSR 10.7, transcriptionally silent in BSF cells, is activated by readthrough transcription upon PP1-1 KD (see below) (Figure 2C). Readthrough transcription was also seen at a termination site between head-to-tail oriented PTUs (Figure 2D).

**Figure 2.**
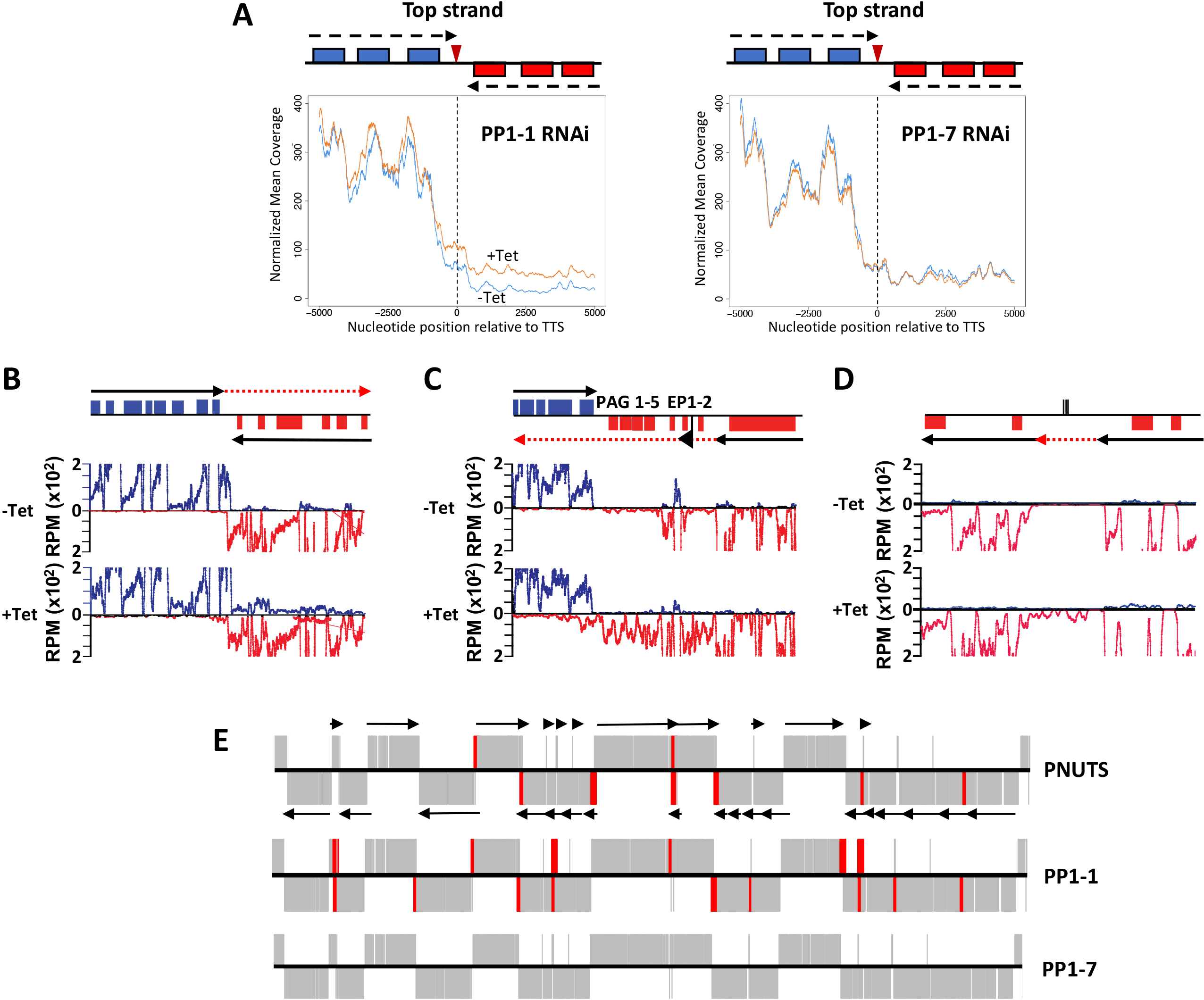
Knockdown of TbPP1-1 results in readthrough at transcription termination sites. (A) Mean top strand coverage at each nucleotide position in the 10 kb surrounding the transcription termination site (TTS) at 39 cSSRs for the PP1-1 and PP1-7 RNAi cell line (as indicated). Uninduced (blue line) or induced (orange line). The schematic represents the protein-coding genes associated with each strand at an “average” convergent TTS and arrows represent the direction of transcription. Plots are orientated that transcription proceeds from the left and terminates at “0”, with the top strand being the coding strand on the left side of the TTS. (B-D) Coverage plots of representative genomic regions. Top; map of the region where ORFs are represented as boxes with the top strand in blue and the bottom strand in red. Black arrows indicate the direction of transcription in each PTU. The red arrow indicates read through transcription following the loss of PP1-1. Bottom; Strand-specific mRNA-seq reads from the control (-Tet) and PP1-1-depleted samples (+Tet) are mapped. Reads that mapped to the top strand are shown in blue and reads that mapped to the bottom strand in red. (B) A region on chromosome 10 from 920-960 kb where J (46), PNUTS (52) and PP1 regulate transcription termination at a cSSR (cSSR 22.3) is shown. See (46) for SSR nomenclature details. (C) Coverage plot of the EP1 locus at cSSR 10.7 (1790-1810 kb). The Pol I promoter is indicated by the flag. (D) A region on chromosome 11 (3443-3463 kb) representing a termination site between head-to-tail (unidirectional) PTUs. The lines on the top strand indicate clustered tRNA genes. (E) Genetic depletion of PP1-1, but not PP1-7, causes transcriptional changes similar to those seen in cells depleted of PNUTS. Gene map of the core section of chromosome 10 is shown. mRNA coding genes on the top strand are indicated by black lines in the top half of the panel, bottom strand by a line in the bottom half. Genes on the top strand are transcribed from left to right and those on the bottom strand are transcribed from right to left. Arrows indicate the direction of transcription of PTUs. Genes that were upregulated >3-fold upon ablation of the indicated factor are highlighted in red. PNUTS RNAi data are from (52).

Differential expression analysis (DESeq2 module) revealed 593 mRNA genes had significantly higher mRNA abundance using a cut-off value of a 3-fold increase and an adjusted *p_adj_*_-_value of ≤ 0.001, in the PP1-1 KD compared to the control (Supplementary Table S1). In contrast, only four transcripts were downregulated using the same threshold. Of the upregulated genes, a majority represent transcripts from genomic loci that are silent under physiological conditions, such as at the end of PTUs in the core of the chromosome or telomeric VSG loci (Table S1). The RNA-seq results confirm our initial RT-PCR analysis of nascent and steady-state RNA indicating a role of PP1-1 regulating Pol II termination and expression of downstream genes of the PTUs located in the chromosome core. The upregulated genes in the PP1-1 mutant located at the 3’-end of PTUs, including the genes (5.3990 and 5.4000) analyzed in Figure 1, map to regions downstream of base J and H3V. The genome-wide mapping to the chromosome core, indicates genes with >3-fold upregulation in the PP1-1 KD were located at regions flanking PTUs, similar to those affected in the PNUTS KD (52) (Figure 2E) and the base J/H3V mutant (46,47). Therefore, consistent with our analysis of the J/H3V and PJW complex mutants, we observe similar increases in expression of genes downstream of termination sites in the PP1-1 mutant, which we previously demonstrated is caused by a defect in Pol II transcription termination resulting in readthrough transcription (46,47,52). In contrast, in the PP1-7 KD only 15 mRNA genes were upregulated using the same threshold, primarily representing silent telomeric and subtelomeric VSG genes (Table S2) (see below). We also noticed some degree of PP1-1 KD in the PP1-7 RNAi (Table S2) that may explain these small changes in gene expression. Consistent with the RT-PCR analysis (Figure 1), no significant termination defects are detected in the chromosome core in the PP1-7 mutant (Figure 2E). All together these results suggest that TbPP1-1, possibly functioning in the PJW complex, is essential for Pol II termination and where loss of function leads to pervasive Pol II transcription of the chromosome core.

### PP1-1 controls transcription of VSG genes from telomeric ESs and subtelomeric arrays

*T. brucei* chromosomes are composed of the transcribed core that contain the primarily Pol II transcribed PTUs and subtelomeric regions that lack a promoter and are thought to be non-expressed. Furthermore, several telomeric regions contain Pol I transcribed PTUs involved in antigenic variation (so called BSF VSG expression sites, BESs) (see Figure 3A). Monoallelic expression of a VSG ES leads to the expression of a single VSG on the surface of the parasite, a key aspect of the strategy BSF cells use to evade the host immune system. In addition to Pol II termination sites distributed within the chromosome core, H3V and base J localize within the silent telomeric BESs (38,41). The loss of H3V/base J or PNUTS KD leads to increased expression of VSG genes from silent telomeric BESs (47,49,52,57). This effect was attributed to the proposed role of the PJW complex in attenuating transcription elongating Pol I within the silent VSG BESs (52). In BSFs, transcription of the silent ESs has been shown to initiate at the Pol I promoter but terminate before the VSG gene (59,60). Consistent with the initial RT-PCR analysis (Figure 1), RNA-seq analysis showed that all VSGs and some ESAGs from silent ESs were significantly upregulated in the PP1-1 KD (Figure 3C; see also Table S1). In contrast, very little change was evident in the PP1-7 KD (Figure 3C). Our analysis indicates an increase of RNA-seq reads that map to ESAGs or VSG genes from silent ESs after PP1-1 KD, reflecting transcription of the entire ES (Figure 3D and S8). The differences in the relative increases among the VSG and ESAG mRNAs are likely due to their differential posttranscriptional regulation (60). Several ESs have dual promoters with an additional promoter upstream of ESAG 10 (58) and were also derepressed in the PP1-1 KD (Figure 3D and S8). In contrast, little to no change was measured in the PP1-7 KD (Figure 3D and S9). In addition to the BESs, depletion of PP1-1 leads to increased VSG expression from the silent telomeric metacyclic ES (MES), which are transcribed monocistronically by Pol I in metacyclic form *T. brucei*. Of eight VSGs in *T. brucei* 427 genome assemblies sited <4 kb downstream of a putative MES promoter, seven were upregulated between 6.5- and 18.5-fold after TbPP1-1 knockdown. Additionally, hundreds of VSG genes and pseudogenes that are located at silent subtelomeric arrays were also expressed after PP1-1 KD (Figure 3C and S10). RT-PCR analysis using specific primers confirmed upregulation of subtelomeric and telomeric VSGs (Figure 3B and S11) and indicated that PP1-1, in contrast to PP1-7, regulates expression of telomeric and subtelomeric regions of the genome. Interestingly, PP1-1 KD did not affect expression of VSG G4 located on a minichromosome that cannot be transcribed due to the lack of a functional promoters (Figure S11). Therefore, VSG derepression by PP1-1 KD requires a promoter somewhere on the chromosome. Altogether, the data indicate that PP1-1 KD leads to expression of transcripts from genomic loci that are typically silent under physiological conditions in BSFs, such as SSRs in the chromosome core and subtelomeric and telomeric arrays.

**Figure 3.**
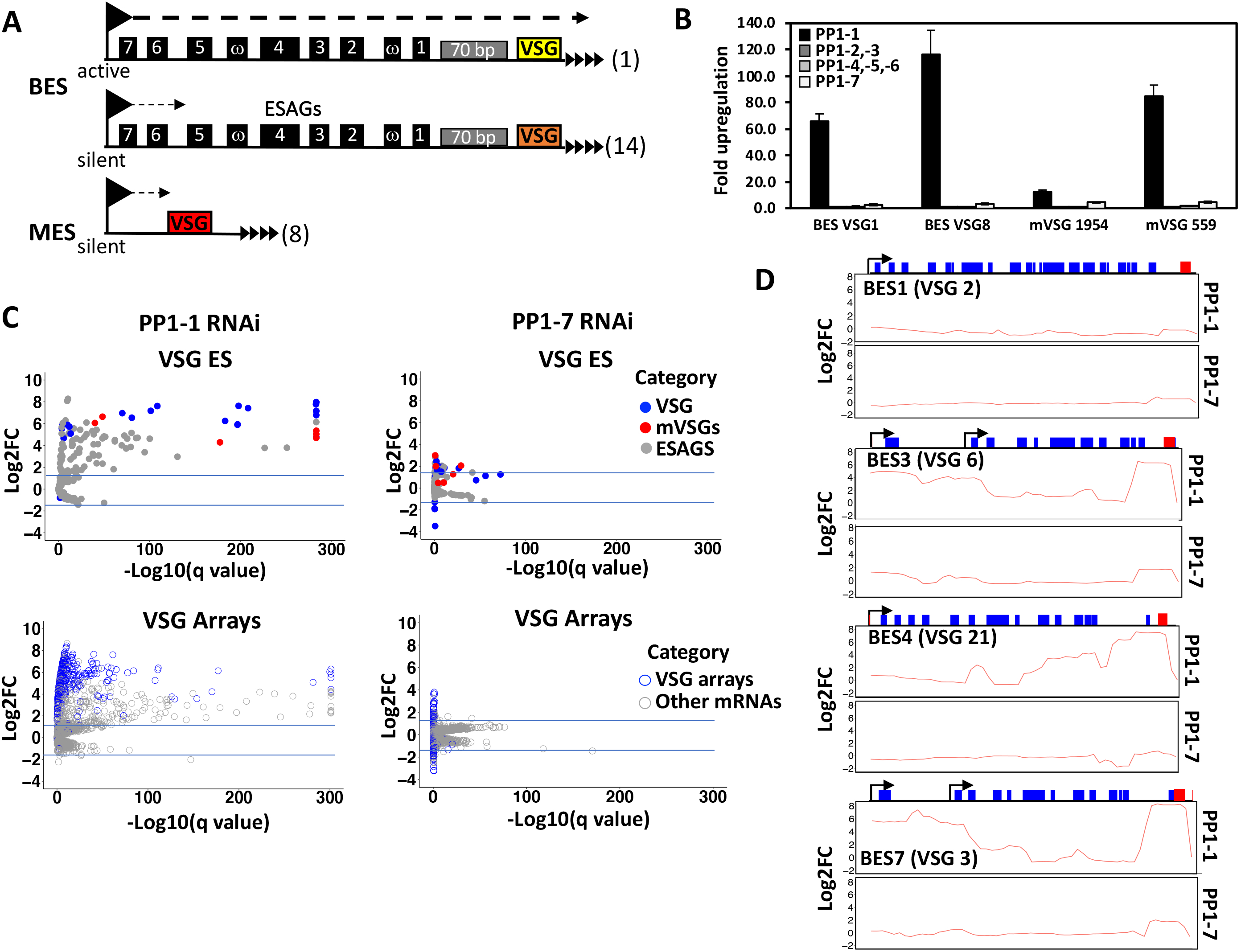
Depletion of PP1-1 disturbs monoalleleic expression of VSG genes. (A) Schematic diagrams of telomeric VSG genome locations; BES, Bloodstream-form Expression sites; and MES, metacyclic expression sites. ESAGs, expression site associated genes. (B) Real-time PCR analysis of VSGs from silent subtelomeric arrays after 48 h of the indicated PP1 isoform knockdown. (C) RNAseq analysis of BFs after 48 h of PP1-1 (left) and PP1-7 (right) knockdown. Top plot shows the VSG and ESAG genes only, and the bottom plot shows VSGs from subtelomeric arrays and mRNAs (Pol II transcribed genes), colored as indicated. See Data Set Table S1 for a list of all transcriptional changes. (D) Comparison of read coverage differences for the top strand at the BESs. RNA-seq reads after 48 h of PP1-1 and PP1-7 knockdown were aligned to the *T. brucei* 427 BES sequences. A diagram of the ES is shown at the top. Promoter (arrow), ESAGs (blue boxes), and VSG (red box) are shown. Fold changes in read coverage comparing – and + Tet were plotted over each BES (Log2 FC). Four of the BES are shown here. See S8 and S9 Figures for all BES. Diagram indicates annotated genes, boxes, within the BES PTU. ESAGs are indicated in blue. The last gene on the right is the VSG gene (red). Promoters are indicated by arrows. Some BESs have two promoters.

### Readthrough transcription leads to transcription of silent Pol I loci by Pol II

Bloodstream form *T. brucei* utilizes RNA Pol I to transcribe rRNA within the nucleolus, as well as the active BES VSG in the ESB (48–50). Metacyclic VSG is expressed by Pol I but only in the insect salivary gland. Procyclin genes (producing abundant surface proteins) are also Pol I transcribed from non-telomeric loci but normally only in insect mid-gut stage cells. The use of Pol I to transcribe certain mRNAs is a unique feature of trypanosomes. We show that various Pol I and Pol II transcribed loci are upregulated upon ablation of PP1-1. Analyzing the extension of nascent RNA at termination sites upon the depletion of epigenetic factors (base J and Histones) (47,48), structural components of the PJW/PP1 complex (52), and depletion of PP1-1 here, suggests the changes in gene expression are due to readthrough transcription by the corresponding polymerase, rather than selective protection of existing RNA from degradation. Therefore, the readthrough transcription defects at the end of Pol II transcribed PTUs in the chromosome core, sometimes leading to de-repression of silent genes immediately downstream of the termination site, are due to readthrough transcription by Pol II. Run-on experiments in the *T. brucei* base J mutant, following Pol II enzyme activity along the chromosome, supports this idea (46). It has been shown that Pol I initiation does occur on silent BES promoters but terminates early preventing transcription of the VSG gene. It was therefore predicted that de-repression of silent VSG genes at the end of the telomeric VSG ES in the PP1-1 KD is because Pol I termination defects within the PTU. However, transcriptome analysis of the subtelomeric VSG arrays (with no identified promoter and presumably never transcribed) suggest that transcription of these silent arrays in the PP1-1 KD could be due to a Pol II termination defect from an adjacent PTU in the chromosome core (Figure S10). Therefore, another possibility is that all gene de-repression events in the PP1-1 mutant are solely due to defects in Pol II termination and corresponding resulting pervasive transcription. Thus, de-repression of the Pol I transcribed telomeric VSGs and subtelomeric VSG arrays would be explained by readthrough transcription of Pol II from the chromosome core. This shift in polymerase transcription due to Pol II termination defects would also explain the upregulation of the procyclin (EP/PAG1) expression site (Figure 2C). Procyclins are surface glycoproteins expressed in procyclic forms of *T. brucei*. Similar to VSG gene transcription, procyclin genes are transcribed by Pol I in a polycistronic unit containing other, coordinately regulated genes. The EP/PAG1 procyclin expression site on chromosome 10 (Figure 4A) includes two copies of procyclin genes followed by a procyclin associated gene (PAG) and is immediately downstream of a Pol II transcribed PTU. According to our model, de-repression of the procyclin loci following PP1 KD is due to Pol II readthrough from the Pol II PTU into the downstream EP/PAG1 procyclin expression site. To test this possibility and determine which RNA polymerase is responsible for the increased transcription of the procyclin locus we performed an RT-PCR assays in PP1-1 ablated cells treated with the Pol I inhibitor BMH-21 for 15 min as described (78). First, we confirmed that Pol I transcription of the rDNA and VSG ES in BS form cells was selectively inhibited by BMH-21. Incubation with 1μM BMH-21, resulted in a dramatic reduction in pre-rRNA and VSG2 precursor transcripts within 15 min, without affecting Pol II derived precursor transcripts from the tubulin array (Figure 4B). In BS form TbPP1-1 RNAi cells, we found that BMH-21 treatment had no effect on the levels of de-repressed nascent RNA from the EP/PAG1 locus (Figure 4C). For example, the EP1 level increased 4-fold in induced PP1-1 RNAi cells before the BMH-21 treatment with no detectable change after treatment (Figure 4C). Using the RNA signal level in induced PP1-1 RNAi as a reference, there was ∼100% of EP1 left after cells were treatment with BMH-21. Similar lack of BMH-21 effect were seen for several precursor RNAs in the EP/PAG1 locus (Figure 4C). In contrast, inhibition of Pol I by BMH-21 in procyclic cells leads to dramatic reduction in RNA from the locus (Figure 4B). As an additional control, we show that block of all transcription using actinomycin D leads to drastic reduction of all RNA measured, including RNAs from the EP/PAG1 locus (Figure 4D). The finding that the RNA level from the EP/PAG locus is insensitive to the Pol I inhibitor BMH-21 in PP1-1 depleted cells, strongly suggests that the majority of RNA from this locus is now transcribed by Pol II resulting from readthrough transcription of the upstream adjacent PTU. Importantly, this ‘polymerase switch’ (expression of Pol I unit by Pol II) supports the model that increased transcripts downstream of termination sites, and gene depression, in the PP1 mutant are primarily due to defects in Pol II termination rather than some sort of selective protection of existing RNA from degradation.

**Figure 4.**
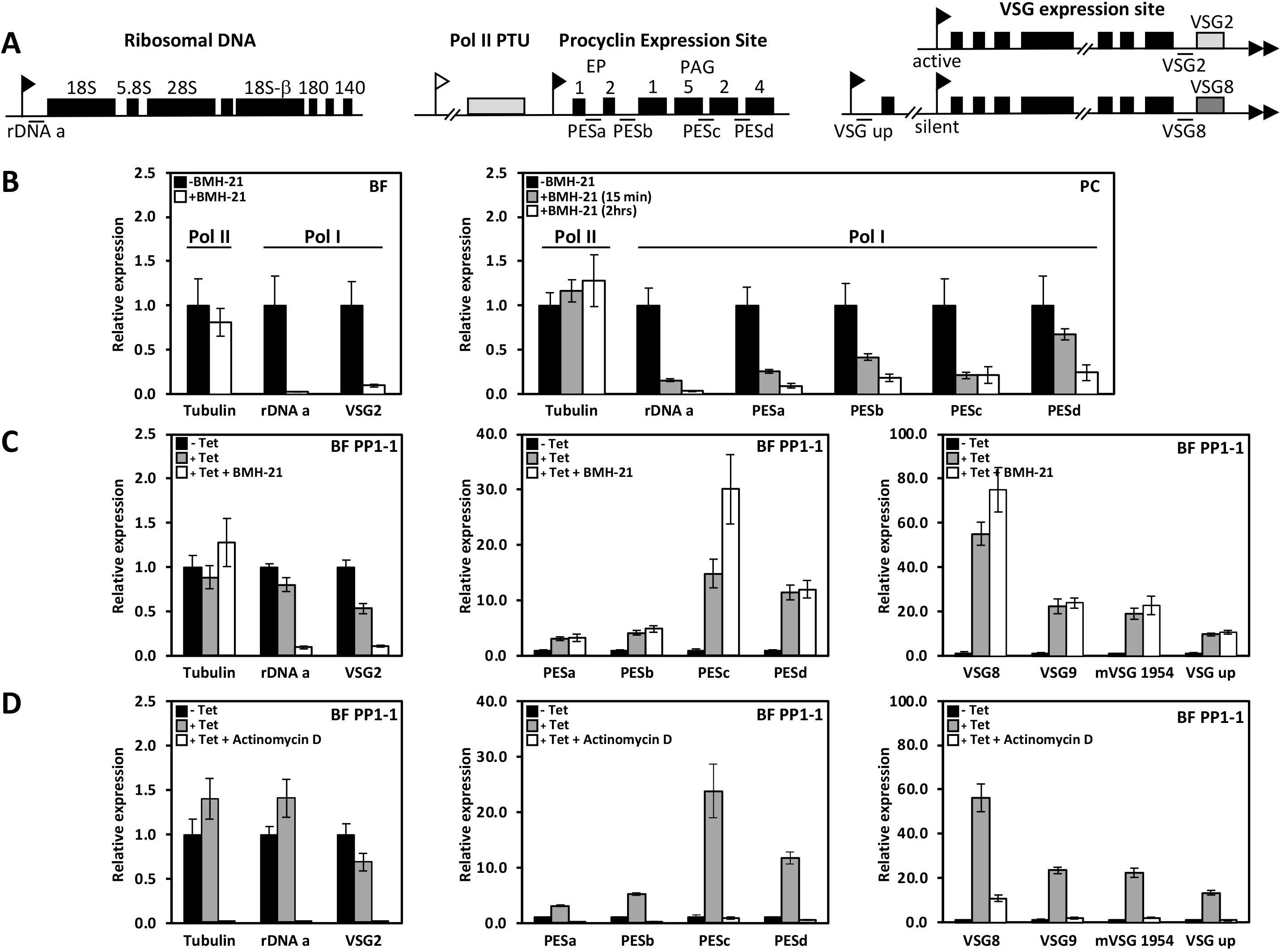
Pol II readthrough transcription leads to Pol II transcription of Pol I loci. (A) Schematic of various Pol I transcription units in T. brucei. Pol I promoters (ribosomal DNA promoter and VSG ES promoter) are indicated with a black flag and Pol II promoter with a white flag. Genes are indicated by boxes. The grey box represents the final gene in the Pol II transcribed PTU immediately upstream of the procyclin locus. qPCR primers are indicated by the line and are labelled. These transcription units are not drawn to scale. (B) Rapid and specific inhibition of Pol I transcription in the presence of BMH-21. BSF (left) *T. brucei* was incubated with 1 μM BMH-21 for 15 minutes. RNA precursor transcripts analyzed were either from the Pol I transcribed rDNA or the active VSG2 precursor transcript from the active VSG ES. In comparison, levels of precursor transcript from the Pol II transcribed alpha-beta tubulin locus remained unaffected. Similar treatment of PC *T. brucei* (right) with BMH-21 affects the Pol I transcription of the procyclin locus as well as the rDNA locus. Initial levels of each precursor transcript before the addition of BMH-21 was set to one. Results shown are the mean of two biological replicates analyzed in triplicate with standard deviation indicated with error bars. (C) Pol II transcribed readthrough products are insensitive to BMH-21. Levels of the indicated precursor transcripts were measured, as in (B), in TbPP1-1 RNAi cells before and 48 hrs after induction of PP1-1 RNAi with and without the BMH-21 treatment for 15 min. (D) All transcription is sensitive to Actinomycin D. As in (C), only cells were incubated with 5 μg/ml Actinomycin D.

To test if Pol II readthrough explains the de-repression of silent telomeric VSG ESs, we further performed the RT-PCR assay for VSG genes in silent BESs in PP1 depleted cells with and without BMH-21. While the VSG in the active BES, VSG2, is sensitive to BMH-21 (as expected) (Figure 4B and C), the increased expression of silent BES VSG8 and VSG9 and the metacyclic VSG1954 in PP1 depleted cells is insensitive to BMH-21 (Figure 4C). Half of the silent BESs in *T. brucei* have a second Pol I promoter approximately 13 kb upstream from the first (58)(Figure 4A). The region between the two promoters encodes *ESAG10* and a variety of pseudogenes, and was particularly derepressed in the PP1 mutant (Figure 3D and S8). Using PCR primers specific for BES9 (Figure 4A), the increased expression in this region is also insensitive to BMH-21 (Figure 4C). RNA-seq indicated that both strands of BESs are transcribed in the PP1 mutant (Figure S8). To differentiate between transcription of each strand we used strand-specific RT-PCR (ssRT-PCR). Figure S12 shows that synthesis of sense and antisense RNA within the silent BES and MES in the PP1 mutant is insensitive to BMH-21. Insensitivity to the Pol I inhibitor BMH-21 suggests that the increased transcription of both strands of silent telomeric VSG ESs is due to Pol II activity. These data indicate Pol II transcription of silent telomeric Pol I loci extending from 13 kB upstream of the BES Pol I promoter to ∼40 kb downstream in the PP1 depleted cells. Overall, the data suggest defects in Pol II termination following the loss of PP1 in *T. brucei* leads to pervasive transcription extending into silent Pol I loci in the chromosome core and silent Pol I loci at the telomeric ends. Furthermore, antisense RNA synthesis at the telomeres would suggest telomere ability, as suggested in yeast (79), to act as Pol II promoters (see below).

### Depletion of PP1-1 increases TERRA formation from the active ES-adjacent telomere

TERRA has been shown to be a product of Pol I readthrough from the active BES into the downstream telomeric repeats in WT *T. brucei* (64,65). To determine if Pol II transcriptional readthrough in the PP1 mutant extends into the telomeric repeats, we first did Northern blotting to examine the TERRA level in PP1-1 RNAi cells. Tubulin RNA level was detected as a loading control. After the induction for 48 hrs, the TERRA level was significantly (3-fold) higher upon ablation of PP1-1 than the control uninduced cells (Figure 5A). In contrast, there is no significant difference in the PP1-7 KD (Figure 5A). Phosphoimager analysis of the blot indicates that depletion of PP1-1 causes no detectable change in TERRA size (Figure 5B). To allow a precise measurement of the TERRA level, we performed dot blot northern hybridization, which showed that TERRA level increased 5-fold upon depletion of TbPP1-1 (Figure 5C). Depletion of PP1-7 resulted in little to no change in TERRA. As a control, all RNA signals were abolished when samples were treated with RNase (Figure 5C). To address the Pol II specificity of the increased TERRA formation in the PP1-1 KD, we treated cells with BMH-21 for 15 min. While BMH-21 abolished TERRA levels in the control (uninduced) cells, the induced levels of TERRA upon ablation of PP1-1 are insensitive to BMH-21 (Figure 5D).

**Figure 5.**
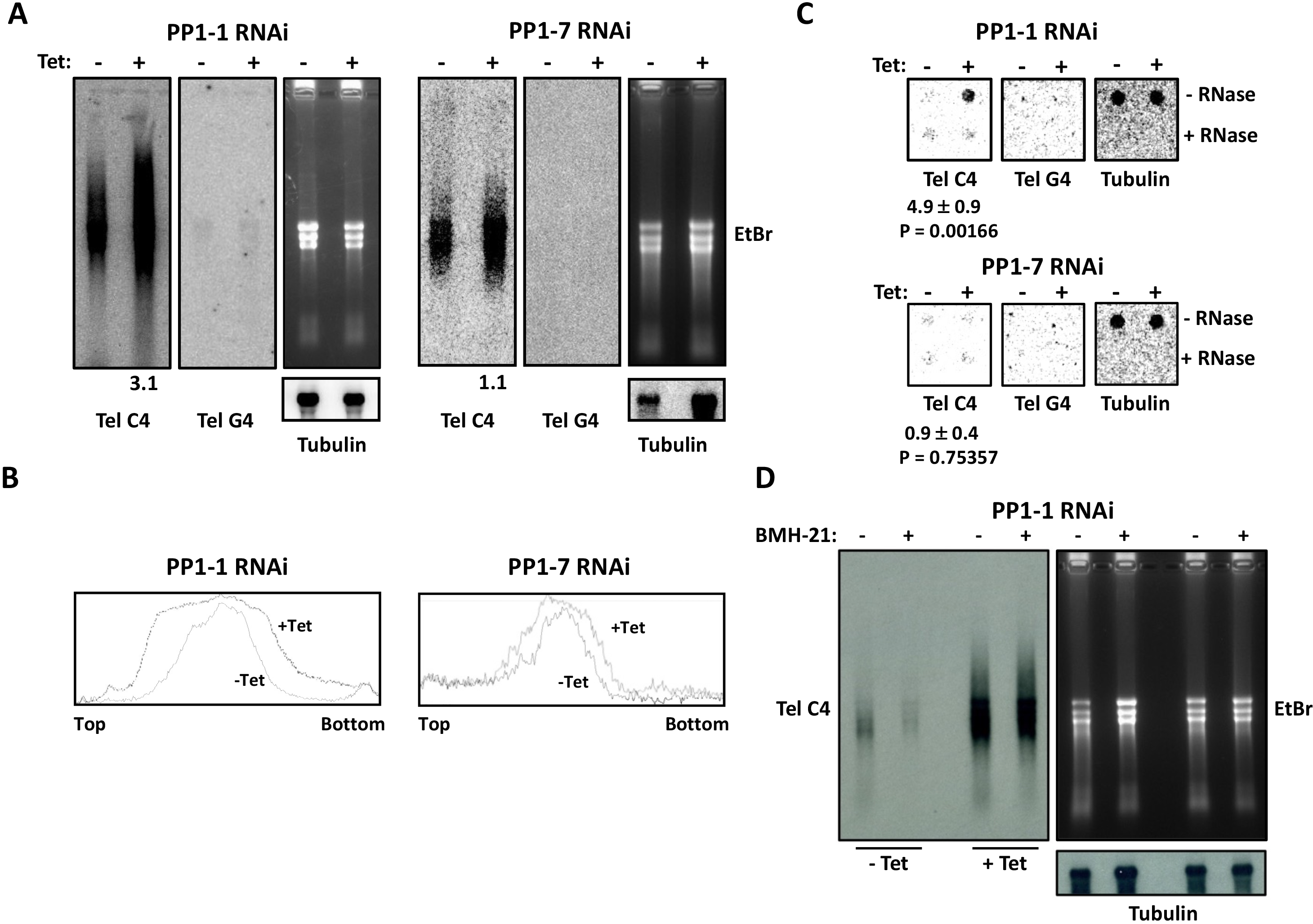
Depletion of TbPP1-1 results in a higher TERRA level. (A) Northern analysis of TERRA in TbPP1-1 (left) and TbPP1-7 (right) RNAi cells. Equal amounts of total RNA was isolated from cells incubated with or without tetracycline for 48 hrs and hybridized with the (CCCTAA)n (TelC4) or the (GGGATT)n (TelG4) probe. Tubulin was detected, and ethidium bromide stained gel, as a loading control. The relative TERRA levels (after normalization against tubulin) were quantitated and indicated at the bottom of the blot. (B) Phosphoimager analysis of the Northern blots in (A). A plot profile of each lane (top to bottom of the blot) is provided. (C) Dot blot northern analysis of TERRA in the PP1-1 and PP1-7 RNAi cell lines before and after adding Tetracycline using a TelC4, TelG4 and tubulin probe. Fold changes in TERRA RNA and tubulin mRNA signal intensities between + and – Tetracycline were calculated from three independent experiments. Standard deviations and P values of unpaired t-test are shown. (D) The stimulated TERRA in the PP1-1 KD is not transcribed by Pol I. Left; a representative TERRA northern blot of samples from PP1-1 RNAi with and without induction with Tetracycline with and without the BMH-21 treatment. Right, ethidium bromide stain of the gel and tubulin hybridization as a loading control.

In WT cells, TERRA is exclusively expressed from the telomere of the active Pol I transcribed VSG ES but not from repeats in the silent VSG ESs (65). To determine the TERRA origin in PP1-1 RNAi cells, we performed the telomere strand-specific RT-PCR as previously described (65,80). After using the CCCTAA repeat containing TELC20 oligo as a primer in the reverse direction, we used primers specific for the active VSG2 or silent VSG3, VSG8, mVSG1954 and mVSG653 genes for PCR analysis. In uninduced PP1-1 RNAi cells, only VSG2-specific primers yielded a clear PCR product (Figure 6, center), as previously observed in WT cells (65). Importantly, after depletion of PP1-1 we observed TERRA transcripts from the silent VSG ESs (Figure 6, center). While the already high level of Pol I transcription in the active ES could mask the increase due to Pol II readthrough activity into this region, a slight increase in TERRA transcripts is observed from this telomeric region as well. In contrast, no changes are observed after depletion of PP1-7 (Figure 6, right).

**Figure 6.**
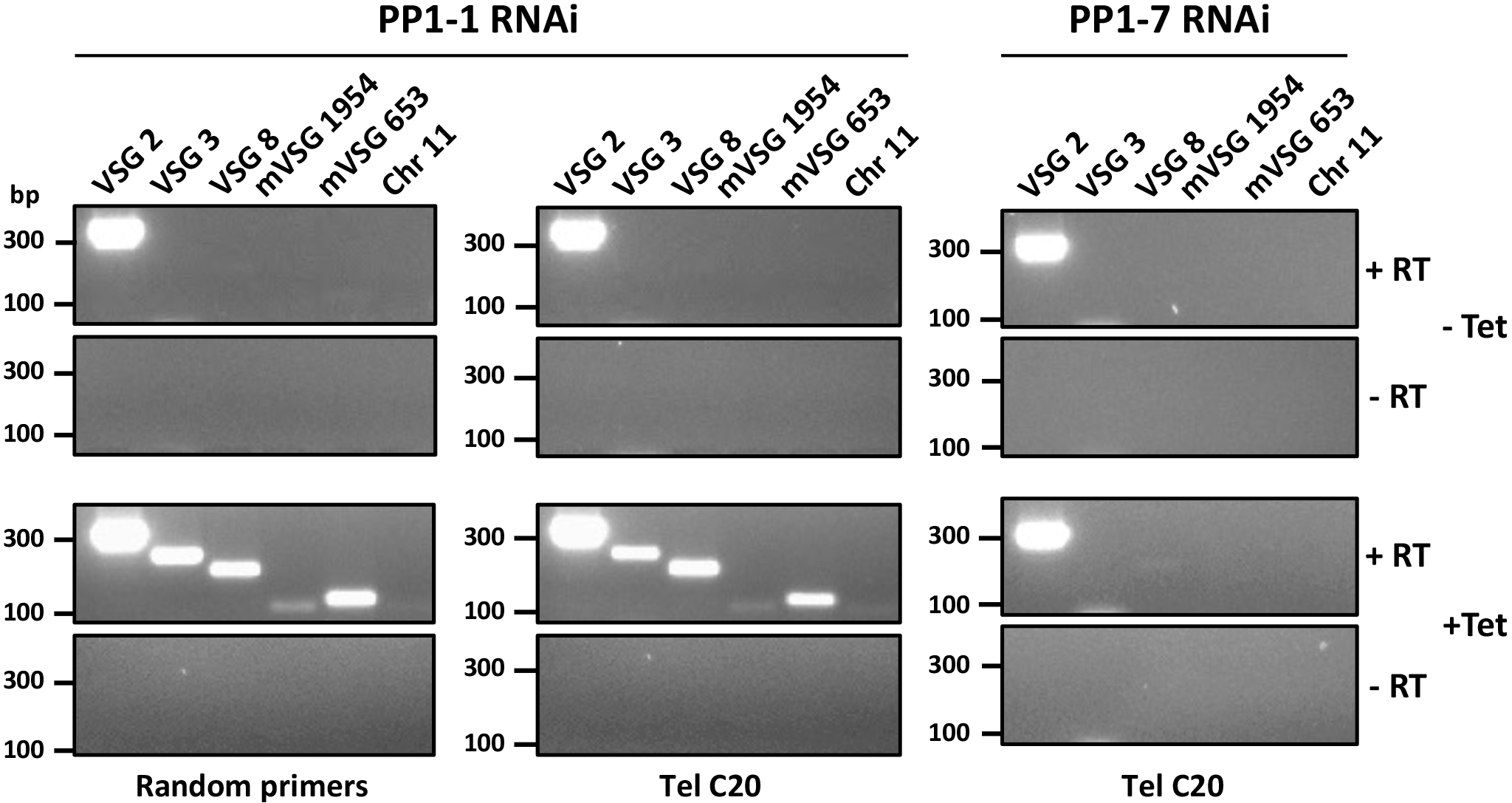
TERRA is transcribed from the active VSG-adjacent telomere in WT and TbPP1-1 depleted cells. Total RNA was purified from the PP1-1 and PP1-7 RNAi cells lines 48 hrs after induction with or without Tetracycline and reverse-transcribed using TelC20 or a random hexamer (as a positive control) as the RT primer (labelled beneath each gel). Reactions without the addition of reverse transcriptase (-RT) are included as a negative control. RT products were PCR amplified using primers specific to the active BES VSG; VSG2, silent BES VSGs; VSG3 and VSG8 (silent), silent MES VSGs; mVSG1954 and mVSG653, and a subtelomeric region of chromosome 11 (Chr 11) (marked on top of each lane), and the PCR products were separated on agarose gels.

Most subtelomeres of *T. brucei* contain VSG gene arrays or VSG ESs, and some contain both (35). As previously described (81), a subtelomere of chromosome 11 lacks any VSG (including subtelomeric VSG arrays and telomeric VSG ES). We identified a unique 100bp sequence within 3kb from the telomere repeat and designed primers that amplify the expected product when using genomic DNA as the template (Figure S13A). Using the TELC20 as the RT primer in the ssRT-PCR assay described above, the chromosome 11 sub-telomere primers confirmed that this VSG-free telomere is not transcribed in WT cells but that deletion of PP1-1 lead to its transcription (Figures 6 and S13B). These results confirm that TERRA is transcribed from the active ES-adjacent telomere but not from silent ES-adjacent telomeres (or other non-ES-adjacent telomeres) in WT cells and suggest that upon depletion of PP1-1 readthrough transcription by Pol II results in transcription of all telomeres.

According to this model, Pol II readthrough results in co-transcription of silent VSGs and TERRA and production of nascent RNA molecules containing both telomeric repeats and the upstream VSG sequence. This nascent RNA is then processed to form, among other RNAs, TERRA (see Figure 7A). To further validate our conclusions regarding TERRA synthesis from previously silent VSG ESs, we performed an RT-PCR experiment to test for this polycistronically generated RNA product. We were aware that under standard denaturing conditions, endogenous priming of telomeric RNA can occur. Furthermore, the regions within the telomeric ES between the VSG gene and telomeric repeats contain GGGTTA-like sequences where alternative (endogenous) priming could lead to false positive results in the telomere strand-specific RT-PCR assay utilized above. To detect the ∼2.3 kb long precursor RNA transcript from the active BES we then performed RT-PCR experiments using the TELC20 primer for RT and different combinations of oligonucleotides for successive PCR reactions (Figure 7A). First, we performed PCR amplifications with 5’VSG2+TELC20. All resulting amplicons are less than 2 kbp, nothing of the expected 2.3 kbp is present in either uninduced or induced PP1-1 RNAi cell lines (Figure 7B). Sequencing indicates the major ∼1.9 kbp product represents mis-priming by the TELC20 oligo ∼300 bp downstream of the VSG2 gene (Figure 7A and S14). Interestingly, the remaining amplicons are produced in PCR reactions with just the TELC20 oligo (Figure 7B), likely representing endogenous priming of RNA produced from telomeric-like sequences distributed throughout the genome. We then performed nested PCR amplification with the 5’VSG2 oligonucleotide + the A1 oligonucleotide that hybridizes to the 3’ end of the active BES prior to the telomeric repeats (Figure 7A). We obtained amplicons of the expected size (∼2.3 kbp) both in induced and uninduced PP1-1 RNAi cell lines, with an increase in the RNA species indicated upon depletion of PP1-1 (Figure 7B). As a control, nested PCR using A3+A1 oligonucleotides obtained the expected 240 bp amplicons representing the 3’ end of the BES. The nested PCR results confirm the existence of TERRA precursor molecules composed of subtelomeric and telomeric sequences in the active BES which increases upon depletion of PP1-1 (Figure 7B). In contrast, similar analysis of the silent VSG3 containing subtelomere does not support the production of TERRA precursors from inactive BESs. RT using TELC20 oligo and PCR with 5’VSG3 +S1 oligonucleotides obtained primarily the non-specific amplicons, but low levels of the product expected from mispriming by TELC20 is present in the PP1-1 depleted cell line (Figure 7C). The low yield prevented attempts for DNA sequencing. Nested PCR amplification with the 5’VSG3+S1 oligonucleotides failed to produce the expected amplicons and PCR with the S3+S1 oligonucleotides led to no detectable products (Figure 7C). These results suggest Pol II transcription in the inactive BES upon PP1-1 depletion is attenuated prior to the telomeric repeats and does not significantly contribute to the increased TERRA population. Mis-priming by the TELC20 oligo in the RT would explain the positive signals from inactive BESs in the telomere strand-specific RT-PCR assay in Figure 6. Therefore, TERRA is transcribed from the active ES-adjacent telomere, but not from silent ES-adjacent telomeres, even upon Pol II readthrough transcription in the PP1-1 mutant cell line.

**Figure 7.**
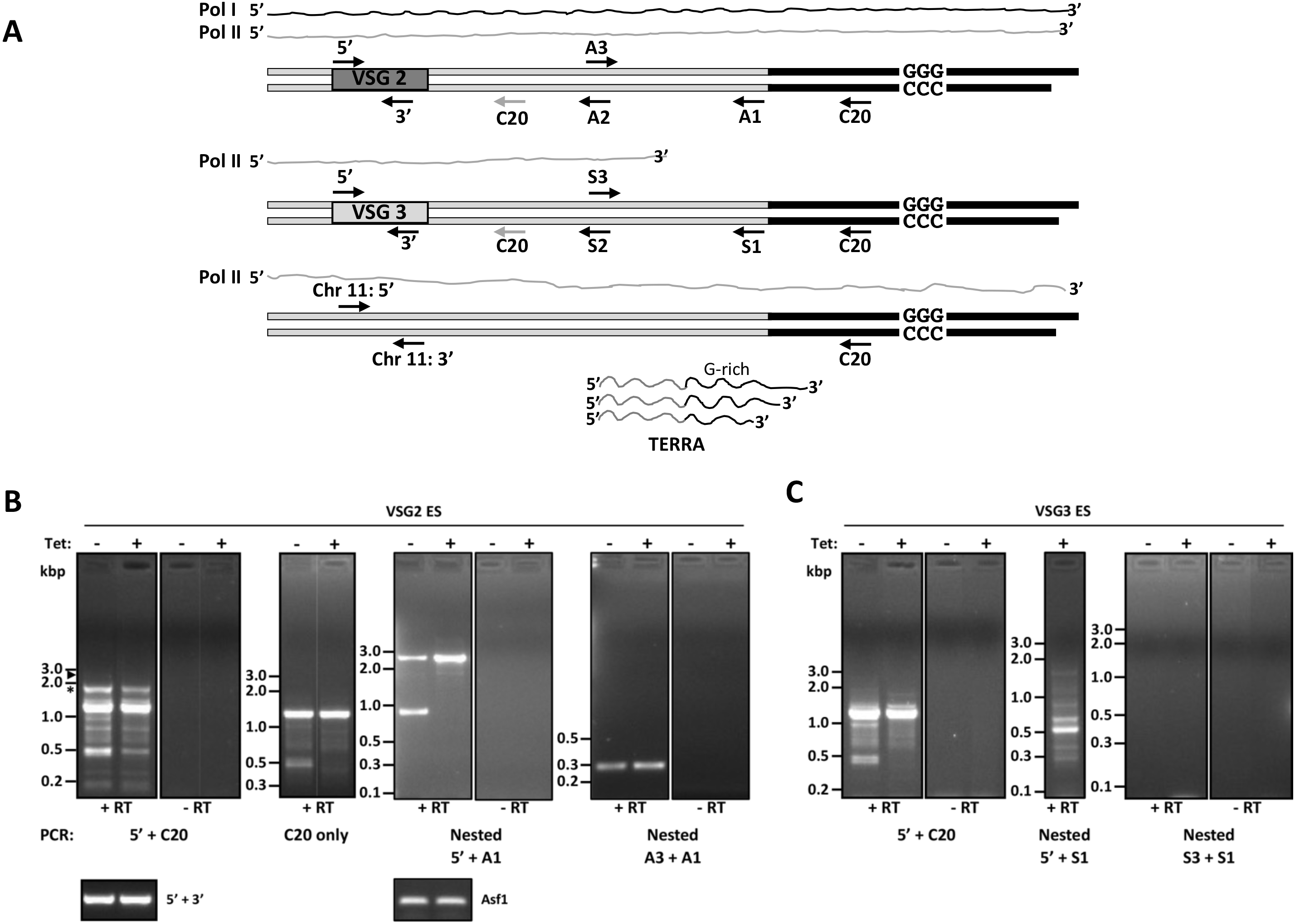
Pol II readthrough transcription of the silent VSG ES is attenuated prior to the telomere repeat. (A) Schematic representation of *T. brucei* chromosome ends and telomeric transcriptome. Telomeric repeats are in black and subtelomeric sequences in white. Black line above represents Pol I transcription of the active BES in WT cells. The grey line represents the proposed extent of Pol II transcription of the subtelomeric loci upon depletion of PP1-1. The large precursor RNA is processed to generate TERRA. Oligonucleotides used for RT-PCR are indicated by arrows pointing in the 5’-3’ direction. The grey arrow for C20 downstream of the VSG gene refers to the mis-priming ability of the primer. The sketch is not in scale. (B) Total RNA from the PP1-1 RNAi cells line 48 hrs after induction with or without tetracycline was reverse-transcribed (+RT) using TelC20 oligonucleotide and cDNA was PCR amplified using VSG2 5’+TelC20 oligonucleotides or just the TelC20 oligonucleotide (left panels). The arrow indicates the expected size product for the long precursor RNA. Asterisk indicates the amplicon that was sequenced shown to represent mis-priming by the TelC20 oligo. Reactions without the addition of reverse transcriptase (-RT) are included as a negative control. Right panels; a portion of this PCR reaction was used in subsequent nested PCR reactions using VSG2 5’+A1 or A3+A1 oligonucleotides. RT-PCR reactions using VSG 5’ and 3’ primers or Asf1 specific primers provides a loading control. Molecular weights are on the left in bp. (C) as in (B), but using primers specific for the silent VSG3 subtelomere.

Overall, the results show that the readthrough phenotype in the PP1-1 KD includes genes that are aberrantly transcribed by Pol II due to upstream termination failure. This includes Pol I transcribed loci that are silent in BS form cells; chromosome internal procyclin expression site, and telomeric VSG expression sites involved in antigenic variation and increased production of TERRA RNA.

### PP1-1 facilitates Pol II-CTD dephosphorylation

PP1-mediated dephosphorylation of the CTD of the largest subunit of the Pol II complex, RPB1, has been shown to underlie the mechanism by which the PTW/PP1 complex modulates transcription termination in human cells. We have recently determined that the Leishmania PJW/PP1 complex similarly regulates transcription termination by dephosphorylation of the RPB1-CTD, by demonstrating PP1-8e phosphatase activity of the complex on Pol II in vitro (53). The identification here of TbPP1-1 as the isotype involved in Pol II transcription led us investigate if it functions via similar mechanisms by assessing whether the depletion of PP1-1 affects Pol II-CTD phosphorylation in vivo. The phosphorylation level of the CTD of Pol II RPB1 was visualized by western blot analysis using anti-HA. In SDS-PAGE, trypanosome RPB1 migrates as a doublet where the upper band is phosphorylated RPB1 and the lower band is the unphosphorylated RPB1 (57,63,72). To confirm the identity of these two forms (where the presence of two forms was due to differences in level of protein phosphorylation) we treated *T. brucei* cells with an inhibitor of PP1, calyculin. We show that treatment with calyculin for 2 hrs is able to convert 100% of Pol II to the phosphorylated species (Figure 8A and B). Therefore, we conclude that the upper and lower bands in our anti-HA western blots represented phosphorylated RPB1 (pRPB1) and dephosphorylated RPB1, respectively. We then utilized the anti-HA western blot to determine the effects of PP1-1 depletion on the status of RPB1 phosphorylation. By western blot, we found that PP1-1 KD, but not PP1-7 KD, increased cellular levels of phosphorylated Pol II while level of total Pol II was unaffected (Figure 8B). We have previously utilized the RNAi of VA protein, and the corresponding significant rapid defect in *T. brucei* cell growth, as a negative control demonstrating defects in Pol II termination were not due to indirect effects of dying cells (52). Thus, as an additional negative control we show that KD of the VA protein, and the associated growth defects, do not lead to changes in Pol II phosphorylation (Figure 8B). These results indicate the defects in Pol II transcription upon depletion of PP1-1 are linked to dephosphorylation of the Pol II-CTD in *T. brucei*.

**Figure 8.**
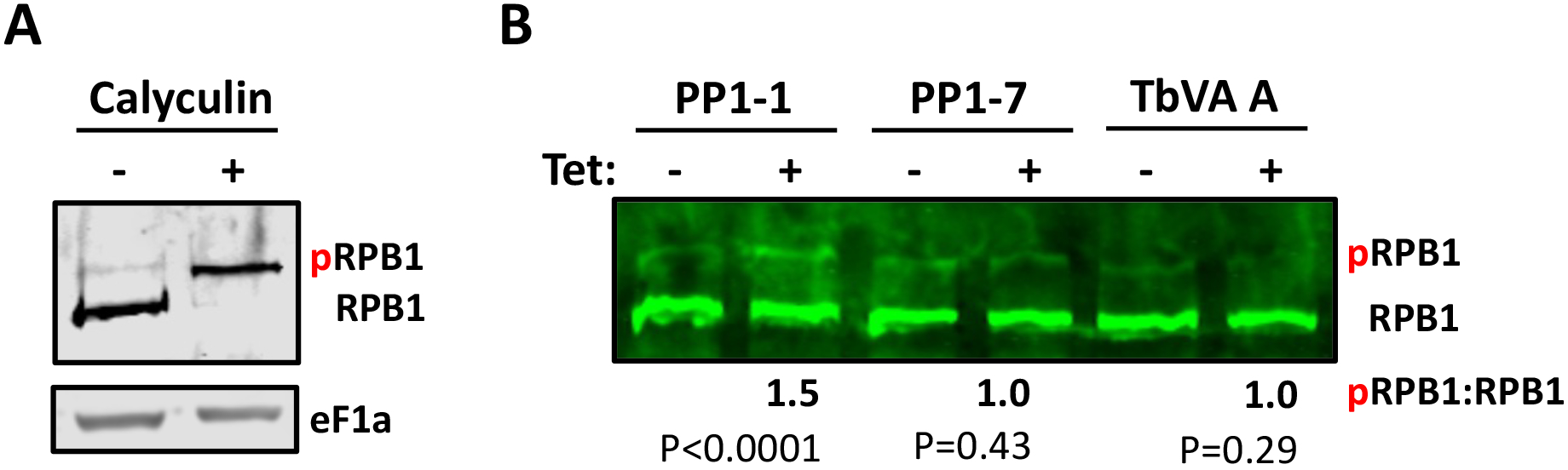
PP1-1 facilitates Pol II-CTD dephosphorylation. (A) TbRNA Pol II is phosphorylated. Western blot of parasite lysate from cells expressing RPB1 endogenously tagged with HA-tag grown in the presence and absence of the PP1 phosphatase inhibitor Calyculin. Blots were probed with anti-HA. Phosphorylated (pRPB1) and dephosphorylated (RPB1) forms of Rpb1 are indicated. EF1a provides a loading control. (B) Western blot of lysate from the indicated RNAi cell lines 24 hrs after induction with or without tetracycline. Ratio of pRPB1/RPB1 from densitometric quantitation of the blots from at least 3 independent experiments is indicated below; with the ratio in untreated cells arbitrarily set to 1. P-values of unpaired t-test are shown.

### PNUTS suppression of Pol II transcription correlates with its PP1-binding activity

In mammalian transcription, PP1 functions through its interaction with PNUTS (as a component of the PTW-PP1 complex), so we wanted to test if this interaction is required for control of Pol II termination in *T. brucei*. Ablation of PNUTS, JBP3 and Wdr82 in *T. brucei* led to similar defects in RNA Pol II transcription termination we describe here for PP1-1 (Figure 2E) (52). This would suggest that PP1-1 functions along with the PNUTS-Wdr82-JBP3 complex in *T. brucei* similar to the activity of LtPP1-8e that we recently demonstrated as part of the PP1-8e-PNUTS-Wdr82-JBP3 complex in Leishmania (53). However, purification of the TbPNUTS-Wdr82-JBP3 complex from *T. brucei* and Co-IP studies, failed to demonstrate stable association of any PP1 isotype, including PP1-1 (52). To address this question, we tested the effect of disrupting the PNUTS-PP1-1 interaction in vivo. Regulatory interactors of PP1 (RIPPOs)(previously referred to as PP1-interacting proteins or PIPs) usually associate with PP1 using a combination of short linear motifs (82–85). The most well-characterized PP1 binding motif of RIPPOs is the RVxF motif that is found in 90% of all RIPPOs including PNUTS, where the second and fourth residues of the motif bury deep in hydrophobic pockets on the PP1 surface, providing an essential stabilizing force. Mutation of hydrophobic positions in the RVxF-binding motif typically abolishes the ability of RIPPO to bind to PP1. We have previously identified the conserved RVxF motif in LtPNUTS and TbPNUTS (52) (Figure S15A) and demonstrated it as an essential PP1 binding motif in Leishmania PNUTS (53). Alanine substitution of the two hydrophobic residues in the RVCW sequence motif of LmPNUTS and LtPNUTS abolished PNUTS-PP1-8e binding in vivo (53,54). RIPPO can also utilize additional motifs beyond the RVxF motif for stable PP1 haloenzyme formation. We have recently employed AlphaFold2 to define an extended RVxF (RVxF-ɸ_R_-ɸɸ) and Phe motif of LtPNUTS involved in binding LtPP1-8e (Figure S15 B and C) (54). Presumably, Leishmania PNUTS utilizing additional motifs beyond the RVxF motif helps stabilize PNUTS:PP1 haloenzyme formation. AlphaFold2 modeling of the TbPNUTS:PP1-1 complex confirms the conserved RVxF motif (RISW) in TbPNUTS, with the two hydrophobic residues of the motif burying deep in hydrophobic pockets on the PP1 surface (Figure S15B). The model also predicts conserved salt bridge interactions between residues that immediately flank the RVXF motif and PP1. Similar interactions occur in the human PNUT-PP1 structure and demonstrated to be critical in LtPP1-8e:PNUTS interaction (54). To elucidate the functional significance of the interaction between PP1 and PNUTS in *T. brucei in vivo*, we examined whether mutation of the conserved hydrophobic residues (I93A-W95A) in the TbPNUTS RVxF motif (converting the RISW motif to RASA) was capable of rescuing the Pol II readthrough transcription phenotypes exhibited by our PNUTS RNAi knock-down (52). To do so, we transfected the PNUTS RNAi cells with a construct allowing Tet regulated expression of an RNAi-resistant C-terminally Flag-tagged wildtype PNUTS (+WT), or the RNAi-resistant Flag-tagged PNUTS with mutations that presumably abrogate PNUTS-PP1 binding (+RASA). As expected, knockdown of PNUTS lead to growth defects and defects in Pol II transcription (including VSG expression), and these effects were rescued by the addition of WT PNUTS (Figure 9). However, the PP1-binding mutant PNUTS (RASA) was unable to rescue the cell growth or transcriptional phenotypes. Analysis of an independent PNUTS RNAi clone, where we expressed a protein A-tagged PNUTS to enhance western blot detection, resulted in similar effects (Figure S16). These data suggest the PNUTS mechanism for controlling Pol II termination in *T. brucei* requires its established interaction with PP1. Therefore, PP1-1 binding to PNUTS is important for its role in Pol II transcription in *T. brucei*.

**Figure 9.**
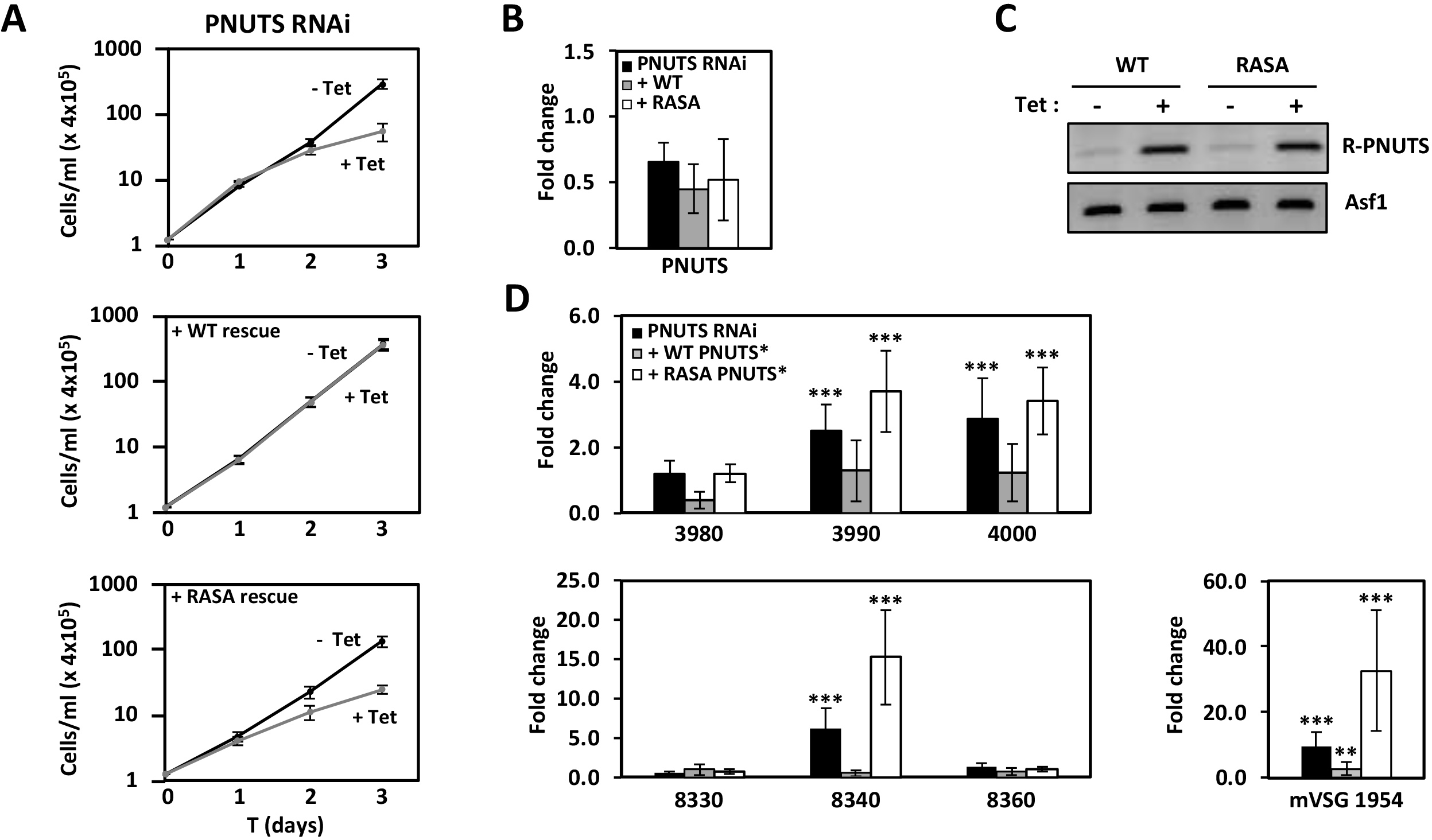
PNUTS suppression of Pol II transcription correlates with its PP1-binding RVXF motif. (A) Top panel; In vitro growth of the PNUTS RNAi cell line with (+Tet) and without (-Tet) induction of RNAi. Bottom panels; growth of PNUTS RNAi cell lines transfected with Tet inducible construct for expression of WT (+WT) or the RVxF mutant (+RASA) of PNUTS recoded to be resistant to RNAi ablation. Growth data are the means (and SEM) of three experiments done in triplicate. (B) qPCR analysis of endogenous PNUTS gene expression 48 hrs after induction of RNAi of the cell lines in (A). (C) PCR analysis of Tet inducible expression of the RNAi resistant WT PNUTS or RASA mutant PNUTS. R-PNUTS; expression of the recoded PNUTS sequence resistant to RNAi. (D) qPCR analysis of Pol II termination defects at chromosome internal Pol II gene clusters as described in Figure 1D and F and at MES VSG (mVSG1954) as in Figure 3B.

## DISCUSSION

### TbPP1-1 functions in termination of Pol II transcription as part of the PJW/PP1 complex

While purification of PNUTS in Leishmania yielded the PJW/PP1 complex, including the sole catalytic component PP1-8e, purification of PNUTS in *T. brucei* only identified the PNUTS-Wdr82-JBP3 structural components. Characterization of PP1-8e function in Leishmania identified its role in Pol II transcription termination; illustrating a conserved PNUTS-PP1 controlled mechanism of Pol II termination among human, yeast and Leishmania. The apparent lack of PP1 stably associated with the Tb PJW complex suggested a unique PP1-independent mechanism in *T. brucei*. Using RT-PCR analysis of nascent RNA and RNA-seq analysis of readthrough RNA genome-wide, we demonstrate here that the depletion of TbPP1-1, in contrast to depletion of the other six functional PP1 isoforms, leads to defects in RNA Pol II transcription termination. The termination defects in the TbPP1-1 KD are similar to defects we measured upon the loss of structural components of the PJW/PP1 complex (TbPNUTS, TbWDR82 and TbJBP3) (52), as well upon the loss of epigenetic marks (base J and H3V) (47,48). Supporting the role of PP1-1 in Pol II transcription, depletion of PP1-1 led to increased levels of phosphorylated RPB1, the large subunit of Pol II. Therefore, linking the defects in Pol II transcription to dephosphorylation of the Pol II-CTD in *T. brucei*. While the shift in phosphorylation status of Pol II upon PP1-1 depletion may seem low, it is specific and approximates the degree of change expected based on transcription termination biology in trypanosomes. Addressing the function of PP1 in dephosphorylating Pol II-CTD *in vivo* was expected to be challenging in *T. brucei*. The analysis of PP1 activity on Pol II *in vivo* (in mammalian and yeast cells) typically utilize P-site-specific antisera, following 1-2 specific P-sites in the heptad repeats known to be linked with termination. Even with this sensitive assay, the change in phosphorylation status of Pol II following inactivation of PP1 is typically small. Complicating the analysis in trypanosomatids, the CTD of RPB1 is divergent and does not carry the heptad or other repetitive motifs. Although 17 P-sites have been identified in *T. brucei* Pol II, no antisera exist for any of the sites. Furthermore, compared with other eukaryotes where Pol II terminates at the end of every gene, the organization of genes in polycistronic arrays in trypanosomatids limits the number of termination sites and thus, the proportion of total Pol II that will be dephosphorylated in the nucleus by PP1 is expected to be small following 50% depletion of PP1-1. Taken together, these findings, along with the presence of a conserved RVxF PP1-binding motif in TbPNUTS, support the idea that TbPP1-1 functions in Pol II transcription while bound to PNUTS in the PJW/PP1 complex. While it was initially surprising that purification of PNUTS from *T. brucei* cells identified the Wdr82 and JBP3 homologs but not the sole catalytic component, recent findings (see below) may explain the unstable nature of TbPNUTS:PP1-1 association and further support PJW/PP1 complex formation and function in *T. brucei*.

PP1 protein phosphatases are thought to have little intrinsic substrate specificity and activity is regulated *in vivo* in a spatiotemporal manner by interacting with RIPPOs, such as PNUTS (3,7). PNUTS-PP1 complex involved in regulating transcription termination is conserved from mammalian to yeast cells (21, 71, 72). Purification of the PJW/PP1 complex from *L. tarentolae* (52, 53) and PP1 isoform specific co-IP studies (54) identified a specific interaction with the PP1-8e isoform among the eight encoded in the Leishmania genome indicating that PNUTS selectively targets PP1-8e to the complex. Identifying its role in Pol II termination, deletion of PP1-8e in *L. major* led to defects in transcription termination at the 3’ end of PTUs, similar to the phenotypes described here for the TbPP1-1 mutant (53). The ability of the purified Leishmania PJW/PP1 complex to dephosphorylate Pol II was dependent on the association of catalytically active form of PP1-8e (53). In contrast, the active complex was unable to dephosphorylate another Leishmania phosphoprotein. Not only did these results provide the first evidence linking base J and Pol II phosphorylation with transcription termination in trypanosomatids, but also indicated that PNUTS association helps regulate PP1-8e substrate specificity. The predicted LtPNUTS:PP1-8e holoenzyme complex, supported by biochemical studies, revealed that LtPNUTS binds PP1-8e using an extended RVxF-ɸ_R_-ɸɸ-Phe motif used by several other RIPPOs including the human PNUTS:PP1 complex (54). The studies indicated that isoform specificity is due to unique sequences in PP1-8e within the catalytic domain and the N- and C-terminus that may be provide binding pockets for the additional short linear motifs of PNUTS beyond the canonical RVxF motif, further stabilizing the PNUTS-PP1 complex. In particular, two residues within the unique C-term α-helix of PP1-8e (P352 and I360), and residues inserted in the catalytic domain (^113^FG^114^), accommodate the Phe motif (F118) in LtPNUTS (54). The other LtPP1 isoforms that lack the PP1-8e specific sequences and fail to bind LtPNUTS, appear to be unable to bind the additional short linear motifs beyond the canonical RVxF motif. The lack of these unique PP1 sequences required for stable LtPP1-8e:PNUTS binding in all TbPP1 isoforms could therefore explain the lack of stable PP1 association during purification of the PJW/PP1 complex from *T. brucei*. AlphaFold prediction of TbPP1-1:PNUTS structure we show here confirms the RVXF motif of TbPNUTS as well as residues flanking the motif involved in conserved salt bridge interactions with PP1. Similar residues in LtPNUTS, and corresponding predicted interactive residues in PP1-8e, are essential for stable LtPNUTS:PP1-8e interaction (54). Demonstrating that perturbation of the conserved PP1-binding site interdicts TbPNUTS essential activities in transcription termination in vivo supports the essential role of PP1-PNUTS interaction in the mechanisms of Pol II termination in *T. brucei*. The fact that PP1-1 does not copurify, and the predicted RVxF-limited PP1:PNUTS interface, suggests PP1-1 is not tightly associated with PNUTS in *T. brucei*; rather, there may be a dynamic or transient association with the transcription termination complex. In fact, despite the relatively high number of PP1 complexes identified in mammalian cells, the highly dynamic nature of these complexes has hindered their functional characterization as well (86). Similar dynamic or transient association is predicted for poly(A) polymerase and the polyadenylation complex (87,88). We propose that the molecular mechanism involved in Pol II transcription termination is similar between Leishmania and *T. brucei* where the PJW/PP1 complex is recruited to termination sites based on base J/JBP3 binding and PP1 dependent dephosphorylation of Pol II leads to attenuation and dissociation of the Pol II complex from the DNA template.

### Allelic exclusion of VSG genes involves the control of Pol II transcription elongation

Base J was discovered based on its link to VSG ES control in BSF *T. brucei* (89). The finding that silent VSG ESs were uniquely resistant to restriction enzyme digestion, led to the discovery of the hypermodified DNA base and its enrichment within repetitive DNA sequences at the telomeric ends of the trypanosome chromosome; including telomeric repeats, the 70bp repeats within the silent VSG ESs (just upstream of the VSG gene) and 50bp repeats upstream of the Pol I promoter. The selective presence of J within the silent ESs and lack within the single active ES suggested its role in VSG gene repression and control of mono-allelic expression. Its presence upstream of the Pol I promoter suggested an additional role as in insulator, preventing the spread of silencing chromatin upstream into the chromosome core and Pol II PTUs. Once J was found to also localize throughout the chromosome of *T. brucei*, as well as within the *L. tarentolae* and *L. major* genome, flanking the Pol II transcribed PTUs and particularly enriched at termination sites, a conserved role of base J (and H3V) in controlling Pol II transcription was demonstrated (41,45-50). As we describe above, this hypothesis now includes a conserved role of J, via the PJW/PP1 complex, in modulating Pol II phosphorylation. Based on the findings presented here, we also propose that one of the functions of base J in *T. brucei* is to control pervasive Pol II transcription from the chromosome core extending into the subtelomeric regions and help maintain mono-allelic expression of VSG. This base J phenotype would be specific to *T. brucei*. Because of the unique lifecycle of *T. brucei* involving growth in the mammalian bloodstream and need to evade the host immune system, telomeric VSG ESs are unique to this trypanosomatid.

In eukaryotes, Pol I exclusively transcribes the large rRNA gene unit (rDNA) in the nucleolar compartment and mRNA is synthesized by RNA pol II. In *T. brucei*, Pol I also synthesizes a subset of abundant pre-mRNAs and the non-coding RNA TERRA. In fact, *T. brucei* is the only known eukaryote with a multifunctional Pol I: transcribing not only rDNA gene units, but also the telomeric localized VSG ES and telomeric repeats producing TERRA, and procyclin ES localized within the chromosome core surrounded by Pol II transcribed PTUs (90). RT-PCR analysis of nascent RNA and RNA-seq analysis of the TbPP1-1 mutant indicates the readthrough RNA defect extends into previously silent regions of the BS form trypanosome chromosome, including the Pol I transcribed procyclin and VSG ES PTUs. Not only is nascent and mRNA from these regions accumulated following the depletion of PP1-1, but we also demonstrate that this is due to a polymerase switch where transcripts are now produced by Pol II. While the production of transcripts from these regions is sensitive to the Pol I inhibitor BMH-21 when they are actively transcribed by Pol I in the appropriate life-stage, they are insensitive when transcribed following de-repression by PP1-1 depletion. Interestingly, it remains to be fully examined if other factors that have been shown to control VSG silencing, where mutation leads to de-repression of silent VSG ES, may similarly be involved in Pol II transcription. Very few studies have demonstrated that de-repression of the VSG ES in the mutant is actually due to Pol I transcription.

### Pol II transcription of TERRA

The G-rich TERRA is transcribed in bloodstream form *T. brucei* via a readthrough product of Pol I transcription of the active VSG ES (64,65). Sensitivity to the Pol I inhibitor BMH-21 (81), confirms TERRA transcription by Pol I in BS form trypanosomes. Telomere proteins TbTRF and its interacting factor TbRAP1 have been shown to regulate the TERRA levels in *T. brucei* (65,81). While depletion of TbTRF and TbRAP1 increases the TERRA level, TERRA transcription is still restricted to Pol I transcription of the active ES-adjacent telomere (81). It is proposed that TRF/RAP1 helps block excessive readthrough of Pol I into the telomeric repeats from the active ES. Interestingly, RAP1 depletion leads to de-repression of the silent ESs and expression of silent VSGs, without transcription extending into the telomere repeats (65); reflecting the lack of detectable change in the ES chromatin structure upon the loss of TbRAP1 (91). We now show that Pol II termination defects in the PP1-1 mutant lead to readthrough transcription that can extend into the active and silent VSG ESs. In the single active VSG ES, Pol II is able to extend into the telomere leading to increased levels of TERRA. Since there is no detectable change in size of the RNA, the increase TERRA levels in the PP1-1 KD is presumably due to increased transcription of the ES by Pol II rather than the ability of polymerase to extend further along the telomeric repeat, as suggested in the TbTRF and TbRAP1 KD (65,81). While Pol II transcription of the silent VSG ESs results in increased expression (de-repression) of ESAGS and VSGs, the polymerase is apparently attenuated prior to the telomeric repeat. Further studies will address if, similar to the RAP1 depletion mutant, this reflects a lack of change in the silent ES chromatin structure involved in telomeric repression upon the loss of TbPP1-1.

Transcription of TERRA by Pol I is unique to trypanosomes. TERRA is transcribed by Pol II in mammals and yeast (92,93) and in some cases a promoter has been identified adjacent to the telomeric repeats (94). Furthermore, transcriptional analyses of telomeres and subtelomeres of various systems have suggested that Pol II production of sense and antisense telomeric lncRNAs is a common feature. The telomeric transcriptome of fission yeast (S. pombe) is comprised of several species of independent Pol II transcripts deriving from chromosome ends (79,95). This includes ‘sense’ subtelomeric transcripts and TERRA as well as ‘antisense’ transcripts. For example, ARIA is a class of transcripts containing C-rich telomeric repeats where the telomere itself acts as a promoter (79). A ∼1 Kb C-rich telomeric RNA was also detected in *T. brucei* PC forms that is absent in BS forms (64). While the G-rich RNA (TERRA) and antisense subtelomeric RNAs are detected in budding and fission yeast (79,95,96), and Leishmania (97), C-rich telomeric RNA seems to be restricted to *S. pombe* and the PC life-stage of *T. brucei*. We do not detect C-rich telomeric RNA in the BS form *T. brucei* WT or PP1-1 mutant by northern blot analysis. Nevertheless, it remains possible that rapidly degrading, unstable C-rich transcripts are produced also in mammals, budding yeast and BS form *T. brucei*. Our high-throughput transcriptional analysis and RT-PCR analysis of nascent RNA have provided evidence of Pol II dependent generation of sense and antisense RNAs at subtelomeres in the Tb PP1-1 KD. These antisense RNAs are also detected in the PNUTS mutant (52). Whether these antisense species represent Pol II transcription initiation within the telomere or subtelomeric region remains to be seen. Regardless, and perhaps more importantly, it is unclear why these antisense transcripts are only seen in the PP1-1 (or PNUTS) mutant BS form trypanosome. Do they represent new Pol II initiation events, allowed by the altered phosphorylation status of Pol II or DNA accessibility due to readthrough transcription of the sense strand, or are they a result of existing Pol II transcription events that are normally terminated by the PNUTS-PP1 complex and rapidly degraded in WT cells? Understanding the molecular events controlling the generation of antisense subtelomeric RNA is an important area for future studies. More significantly, these observations from the PP1-1 mutant provide powerful tools for future studies into the novel non-coding RNA TERRA.

Overall, this work provides new insight into the molecular mechanism utilized to control Pol II transcription termination at the end of PTUs in these divergent human pathogens. The data support conserved function of proteins involved in transcription termination among eukaryotes, despite the need of trypanosomatids to bypass termination at the 3’ end of every gene and terminate in base J specific manner at the end of long polycistronic gene arrays and prevent the activation of silent Pol I loci.

## DATA AVAILABILITY

All sequencing data discussed in this publication have been deposited in NCBI’s Gene Expression Omnibus and are accessible through GEO Series accession number GSE244975.

## SUPPLEMENTARY DATA

Supplementary Data are available at NAR online: Supplementary Tables S1-S4 and Supplementary Figures S1-S16.

## ACKNOWLEDGEMENTS

We are grateful for Erin Campbell for preparing the mRNA-seq libraries.

Supporting Information Table S1. Genes with changes in mRNA levels after RNAi depletion of PP1-1 and PP1-7. The genes showing the largest (>2log fold change) average (between the three replicates) significant (*p*-value_adj_<0.001) increase in RNA abundance, as determined by RNA-seq analysis, are shown. Gene descriptions (annotation and accession number), logfold upregulation and *p*-value adj are also listed. Level of PP1 transcripts for all isoforms is shown at the bottom.

Supporting Information Table S2. *T. brucei* gene expression changes after depletion of PP1-7 as in Table S1.

Supporting Information Table S3. High-throughput sequencing information. Information about all sequencing data generated in this study is listed.

Supporting Information Table S4. Oligos used in these studies.

