## Supplemental Figures for "Protein Phosphatase PP1 Regulation of Pol II Phosphorylation is Linked to Transcription Termination and Allelic Exclusion of VSG Genes and TERRA in Trypanosomes"

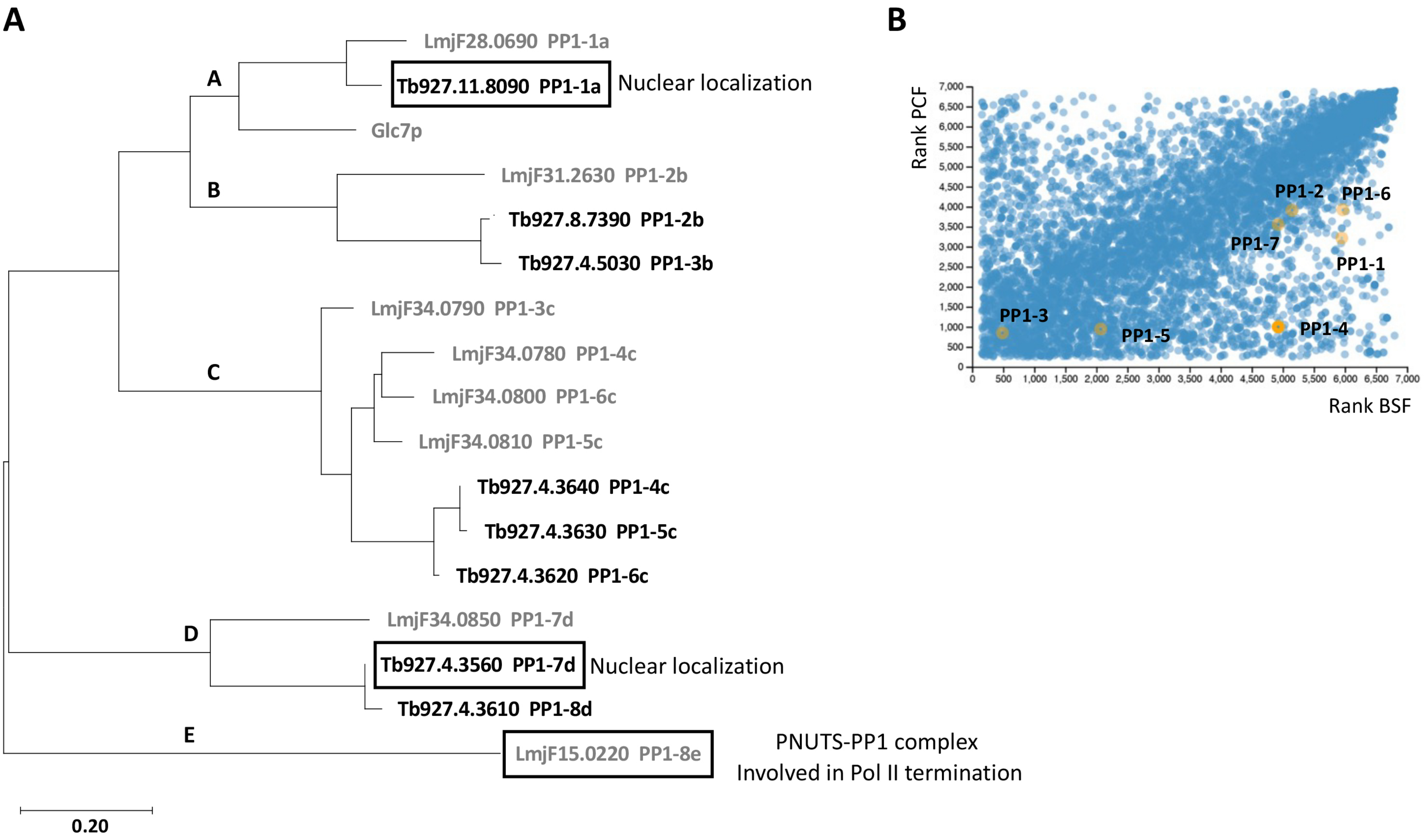
A
LmjF28.0690 PP1-1a
Tb927.11.8090 PP1-1a
Nuclear localization
Glc7p
B
LmjF31.2630 PP1-2b
Tb927.8.7390 PP1-2b
Tb927.4.5030 PP1-3b
C
LmjF34.0790 PP1-3c
LmjF34.0780 PP1-4c
LmjF34.0800 PP1-6c
LmjF34.0810 PP1-5c
Tb927.4.3640 PP1-4c
Tb927.4.3630 PP1-5c
Tb927.4.3620 PP1-6c
D
LmjF34.0850 PP1-7d
Tb927.4.3560 PP1-7d
Nuclear localization
Tb927.4.3610 PP1-8d
E
LmjF15.0220 PP1-8e
PNUTS-PP1 complex
Involved in Pol II termination
0.20
B
Rank PCF
Rank BSF
PP1-1
PP1-2
PP1-3
PP1-4
PP1-5
PP1-6
PP1-7

Figure S1

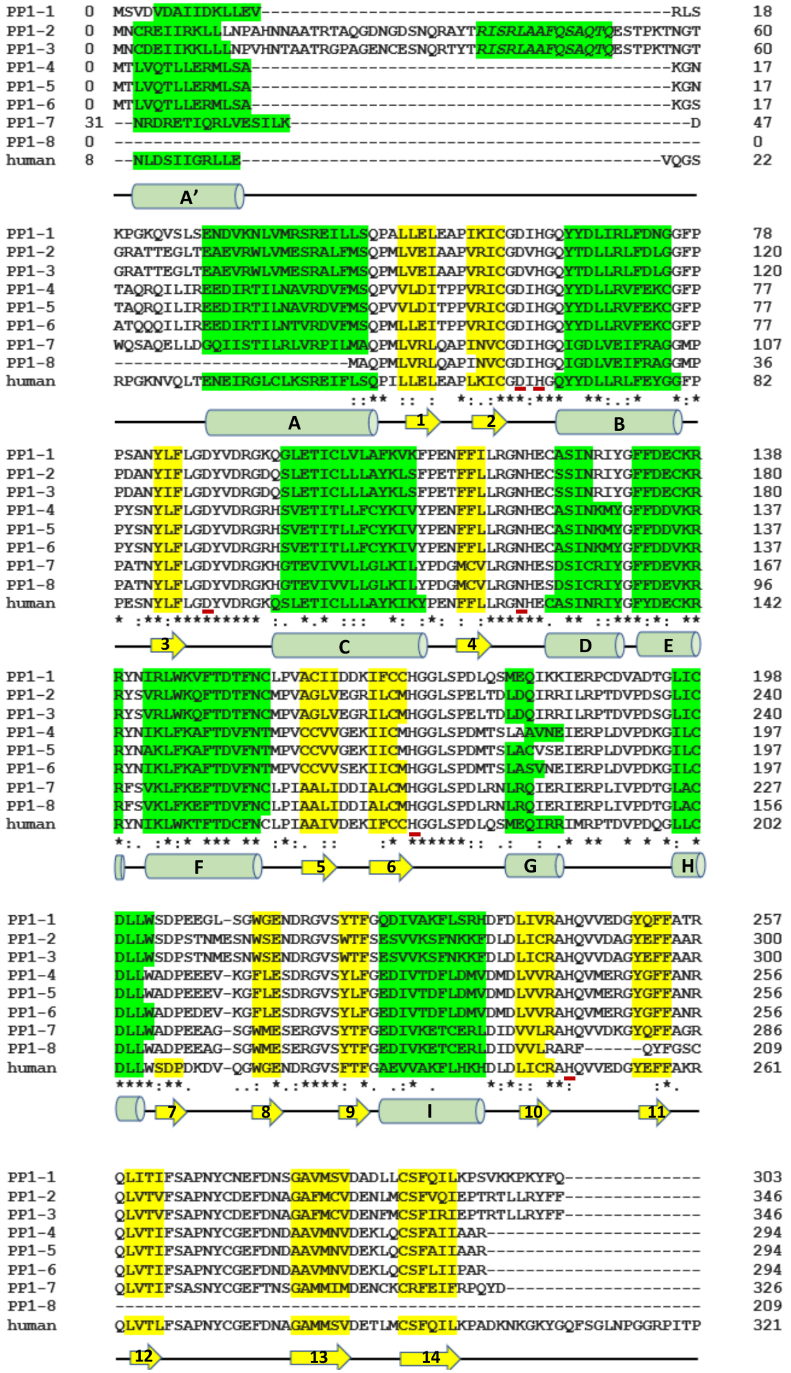

PP1-1 0 MSVDVDAIIDKLLEV------------------------------------------RLS 18
PP1-2 0 MNCREIIRKLLLNPAHNNAATRTAQGDNGDSNQRAYTRISRLAAFQSAQTQESTPKTNGT 60
PP1-3 0 MNCDEIIKKLLLNPVHNTAATRGPAGENCESNQRTYTRISRLAAFQSAQTQESTPKTNGT 60
PP1-4 0 MTLVQTLLERMLSA-------------------------------------------KGN 17
PP1-5 0 MTLVQTLLERMLSA-------------------------------------------KGN 17
PP1-6 0 MTLVQTLLERMLSA-------------------------------------------KGS 17
PP1-7 31 --NRDRETIQRLVESILK-----------------------------------------D 47
PP1-8 0 ------------------------------------------------------------ 0
human 8 --NLDSIIGRLLE-------------------------------------------VQGS 22
A'
PP1-1 KPGKQVSLSENDVKNLVMRSREILLSQPALLELEAPIKICGDIHGQYYDLIRLFDNGGFP 78
PP1-2 GRATTEGLTEAEVRWLVMESRALFMSQPMLVEIAAPVRICGDVHGQYTDLLRLFDLGGFP 120
PP1-3 GRATTEGLTEAEVRWLVMESRALFMSQPMLVEIAAPVRICGDVHGQYTDLLRLFDLGGFP 120
PP1-4 TAQRQILIREEDIRTILNAVRDVFMSQPVVLDITPPVRICGDIHGQYYDLLRVFEKCGFP 77
PP1-5 TAQRQILIREEDIRTILNAVRDVFMSQPVVLDITPPVRICGDIHGQYYDLLRVFEKCGFP 77
PP1-6 ATQQQILIREEDIRTILNTVRDVFMSQPMLLEITPPVRICGDIHGQYYDLLRVFEKCGFP 77
PP1-7 WQSAQELLDGQIISTILRLVRPILMAQPMLVRLQAPINVCGDIHGQIGDLVEIFRAGGMP 107
PP1-8 ------------------------MAQPMLVRLQAPINVCGDIHGQIGDLVEIFRAGGMP 36
human RPGKNVQLTENEIRGLCLKSREIFLSQPILLELEAPLKICGDIHGQYYDLLRLFEYGGFP 82
::** :: : *:.:***:*** **:.:* *:*
A 1 2 B
PP1-1 PSANYLFLGDYVDRGKQGLETICLVLAFKVKFPENFFILRGNHECASINRIYGFFDECKR 138
PP1-2 PDANYIFLGDYVDRGDQSLETICLLLAYKLSFPETFFLLRGNHECSSINRIYGFFDECKR 180
PP1-3 PDANYIFLGDYVDRGDQSLETICLLLAYKLSFPETFFLLRGNHECSSINRIYGFFDECKR 180
PP1-4 PYSNYLFLGDYVDRGRHSVETITLLFCYKIVYPENFFLLRGNHECASINKMYGFFDDVKR 137
PP1-5 PYSNYLFLGDYVDRGRHSVETITLLFCYKIVYPENFFLLRGNHECASINKMYGFFDDVKR 137
PP1-6 PYSNYLFLGDYVDRGRHSVETITLLFCYKIVYPENFFLLRGNHECASINKMYGFFDDVKR 137
PP1-7 PATNYLFLGDYVDRGKHGTEVIVVLLGLKILYPDGMCVLRGNHESDSICRIYGFFDEVKR 167
PP1-8 PATNYLFLGDYVDRGKHGTEVIVVLLGLKILYPDGMCVLRGNHESDSICRIYGFFDEVKR 96
human PESNYLFLGDYVDRGKQSLETICLLLAYKIKYPENFFLLRGNHECASINRIYGFYDECKR 142
* :**:********* :. *.* ::: *: :*: : :******. ** ::***:*: **
3 C 4 D E
PP1-1 RYNIRLWKVFTDTFNCLPVACIIDDKIFCCHGGLSPDLQSMEQIKKIERPCDVADTGLIC 198
PP1-2 RYSVRLWKQFTDTFNCMPVAGLVEGRILCMHGGLSPELTDLDQIRRILRPTDVPDSGLIC 240
PP1-3 RYSVRLWKQFTDTFNCMPVAGLVEGRILCMHGGLSPELTDLDQIRRILRPTDVPDSGLIC 240
PP1-4 RYNIKLFKAFTDVFNTMPVCCVVGEKIICMHGGLSPDMTSLAAVNEIERPLDVPDKGILC 197
PP1-5 RYNAKLFKAFTDVFNTMPVCCVVGEKIICMHGGLSPDMTSLACVSEIERPLDVPDKGILC 197
PP1-6 RYNIKLFKAFTDVFNTMPVCCVVSEKIICMHGGLSPDMTSLASVNEIERPLDVPDKGILC 197
PP1-7 RFSVKLFKEFTDVFNCLPIAALIDDIALCMHGGLSPDLRNLRQIERIERPLIVPDTGLAC 227
PP1-8 RFSVKLFKEFTDVFNCLPIAALIDDIALCMHGGLSPDLRNLRQIERIERPLIVPDTGLAC 156
human RYNIKLWKTFTDCFNCLPIAAIVDEKIFCCHGGLSPDLQSMEQIRRIMRPTDVPDQGLLC 202
*:. :*:* *** ** :*:. :: :* ******:: .: : .* ** * * *: *
F 5 6 G H
PP1-1 DLLWSDPEEGL-SGWGENDRGVSYTFGQDIVAKFLSRHDFDLIVRAHQVVEDGYQFFATR 257
PP1-2 DLLWSDPSTNMESNWSENDRGVSWTFSESVVKSFNKKFDLDLICRAHQVVDAGYEFFAAR 300
PP1-3 DLLWSDPSTNMESNWSENDRGVSWTFSESVVKSFNKKFDLDLICRAHQVVDAGYEFFAAR 300
PP1-4 DLLWADPEEEV-KGFLESDRGVSYLFGEDIVTDFLDMVDMDLVVRAHQVMERGYGFFANR 256
PP1-5 DLLWADPEEEV-KGFLESDRGVSYLFGEDIVTDFLDMVDMDLVVRAHQVMERGYGFFANR 256
PP1-6 DLLWADPEDEV-KGFLESDRGVSYLFGEDIVTDFLDMVDMDLVVRAHQVMERGYGFFANR 256
PP1-7 DLLWADPEEAG-SGWMESERGVSYTFGEDIVKETCERLDIDVVLRAHQVVDKGYQFFAGR 286
PP1-8 DLLWADPEEAG-SGWMESERGVSYTFGEDIVKETCERLDIDVVLRARF------QYFGSC 209
human DLLWSDPDKDV-QGWGENDRGVSFTFGAEVVAKFLHKHDLDLICRAHQVVEDGYEFFAKR 261
****:**. ..: *.:****: *. .:* . *:*:: **: :*.
7 8 9 I 10 11
PP1-1 QLITIFSAPNYCNEFDNSGAVMSVDADLLCSFQILKPSVKKPKYFQ-------------- 303
PP1-2 QLVTVFSAPNYCDEFDNAGAFMCVDENLMCSFVQIEPTRTLLRYFF-------------- 346
PP1-3 QLVTVFSAPNYCDEFDNAGAFMCVDENFMCSFIRIEPTRTLLRYFF-------------- 346
PP1-4 QLVTIFSAPNYCGEFDNDAAVMNVDEKLQCSFAIIAAR---------------------- 294
PP1-5 QLVTIFSAPNYCGEFDNDAAVMNVDEKLQCSFAIIAAR---------------------- 294
PP1-6 QLVTIFSAPNYCGEFDNDAAVMNVDEKLQCSFLIIPAR---------------------- 294
PP1-7 QLVTIFSASNYCGEFTNSGAMMIMDENCKCRFEIFRPQYD-------------------- 326
PP1-8 ------------------------------------------------------------ 209
human QLVTLFSAPNYCGEFDNAGAMMSVDETLMCSFQILKPADKNKGKYGQFSGLNPGGRPITP 321
12 13 14


**Figure S2**

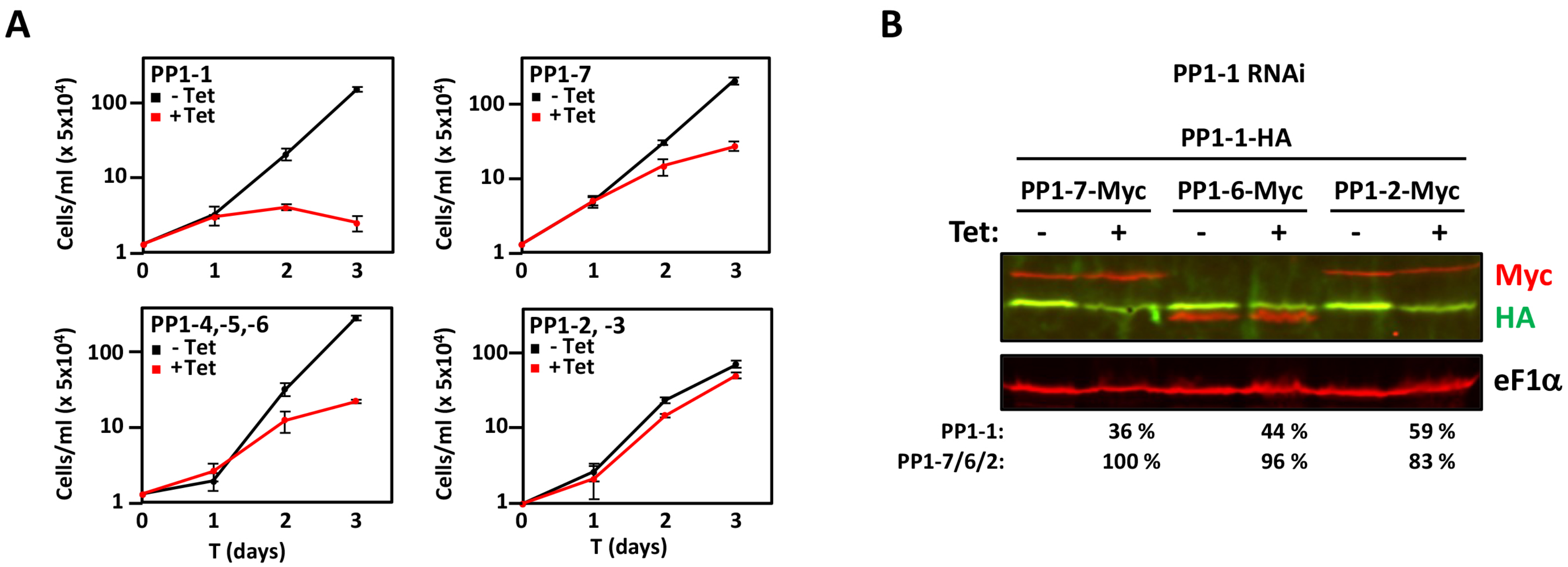
A
PP1-1
- Tet
+Tet
PP1-7
PP1-4,-5,-6
PP1-2, -3
Cells/ml (x 5x10⁴)
100
10
1
0
1
2
3
T (days)
B
PP1-1 RNAi
PP1-1-HA
PP1-7-Myc
PP1-6-Myc
PP1-2-Myc
Tet:
-
+
Myc
HA
eF1α
PP1-1: 36 % 44 % 59 %
PP1-7/6/2: 100 % 96 % 83 %

Figure S3

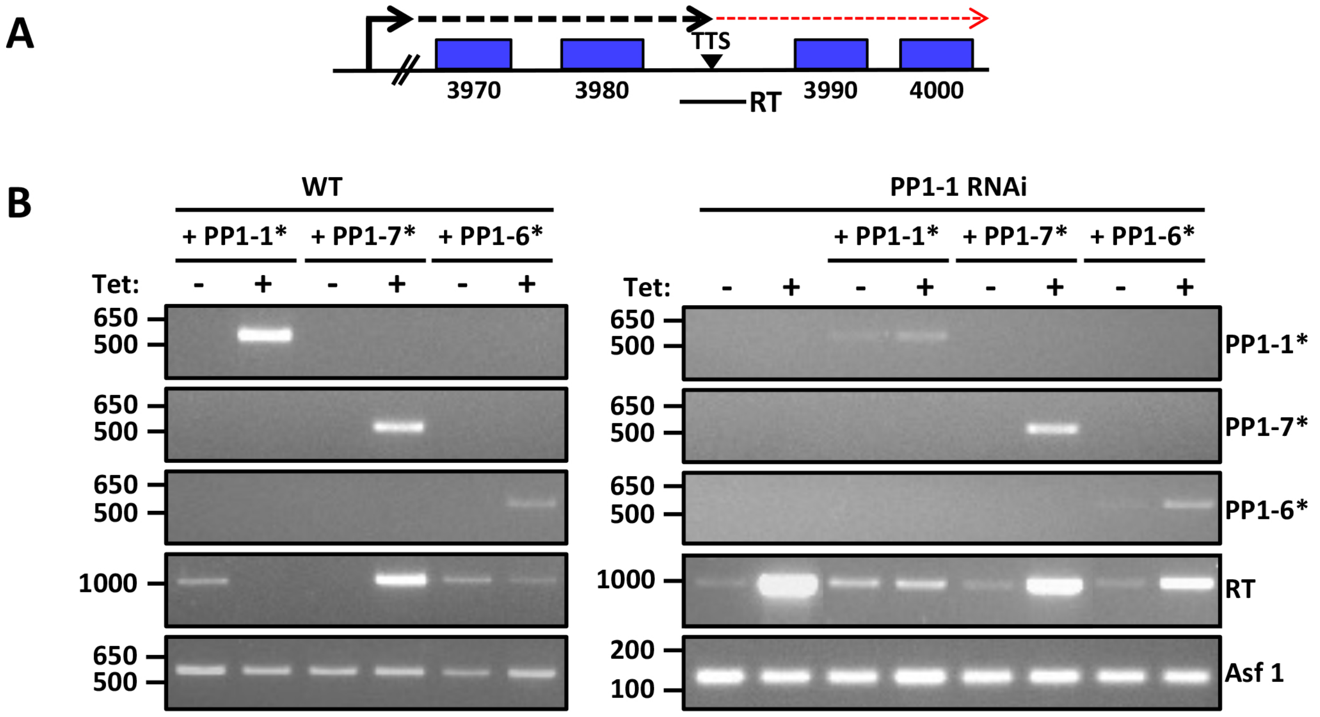
A
TTS
3970
3980
3990
4000
RT
B
WT
PP1-1 RNAi
+ PP1-1*
+ PP1-7*
+ PP1-6*
Tet:
650
500
1000
200
100
PP1-1*
PP1-7*
PP1-6*
RT
Asf 1

Figure S4

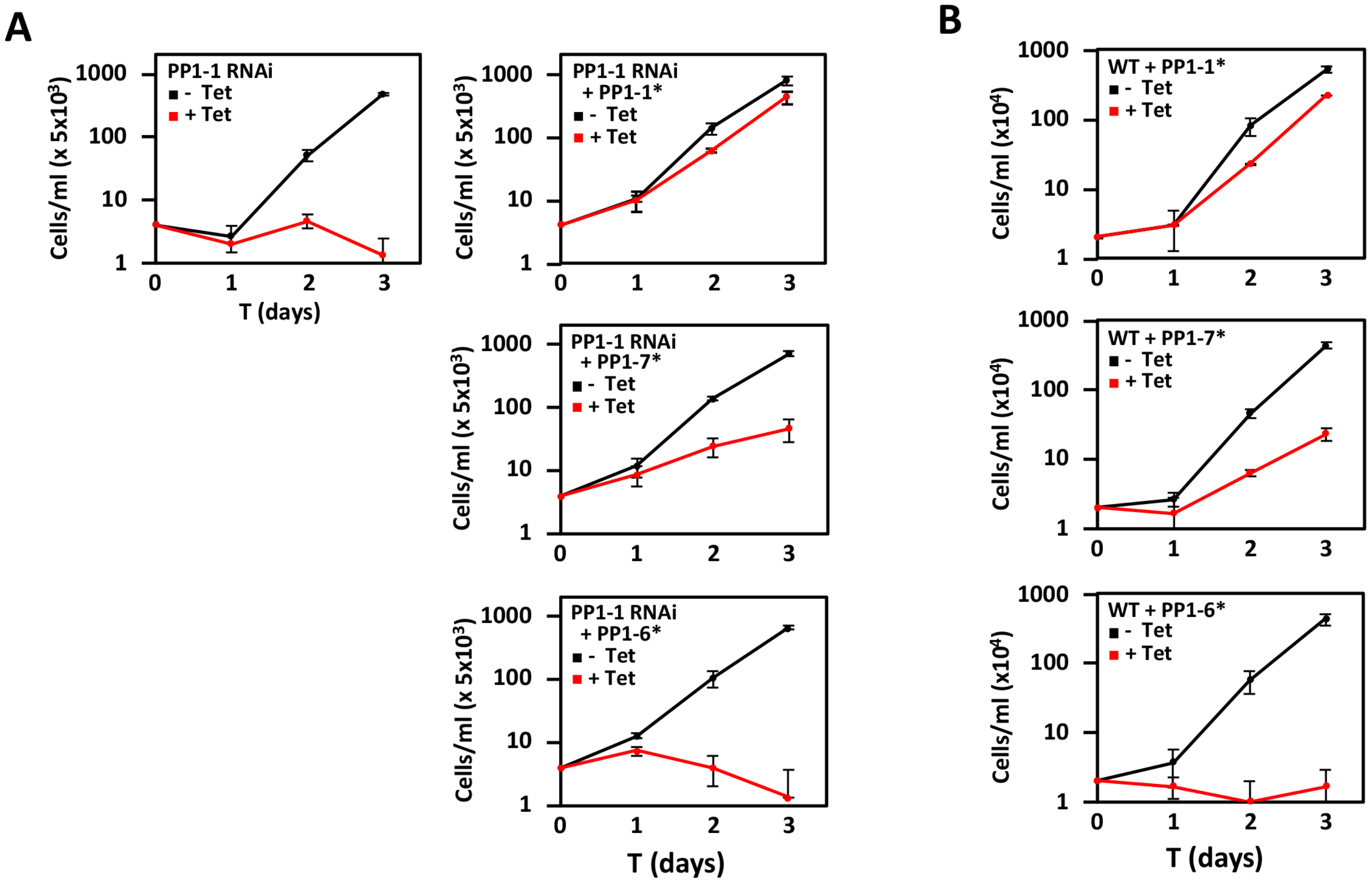
A
B
PP1-1 RNAi
- Tet
+ Tet
PP1-1 RNAi + PP1-1*
PP1-1 RNAi + PP1-7*
PP1-1 RNAi + PP1-6*
WT + PP1-1*
WT + PP1-7*
WT + PP1-6*
Cells/ml (x 5x10³)
Cells/ml (x10⁴)
T (days)
1
10
100
1000
0
1
2
3

Figure S5

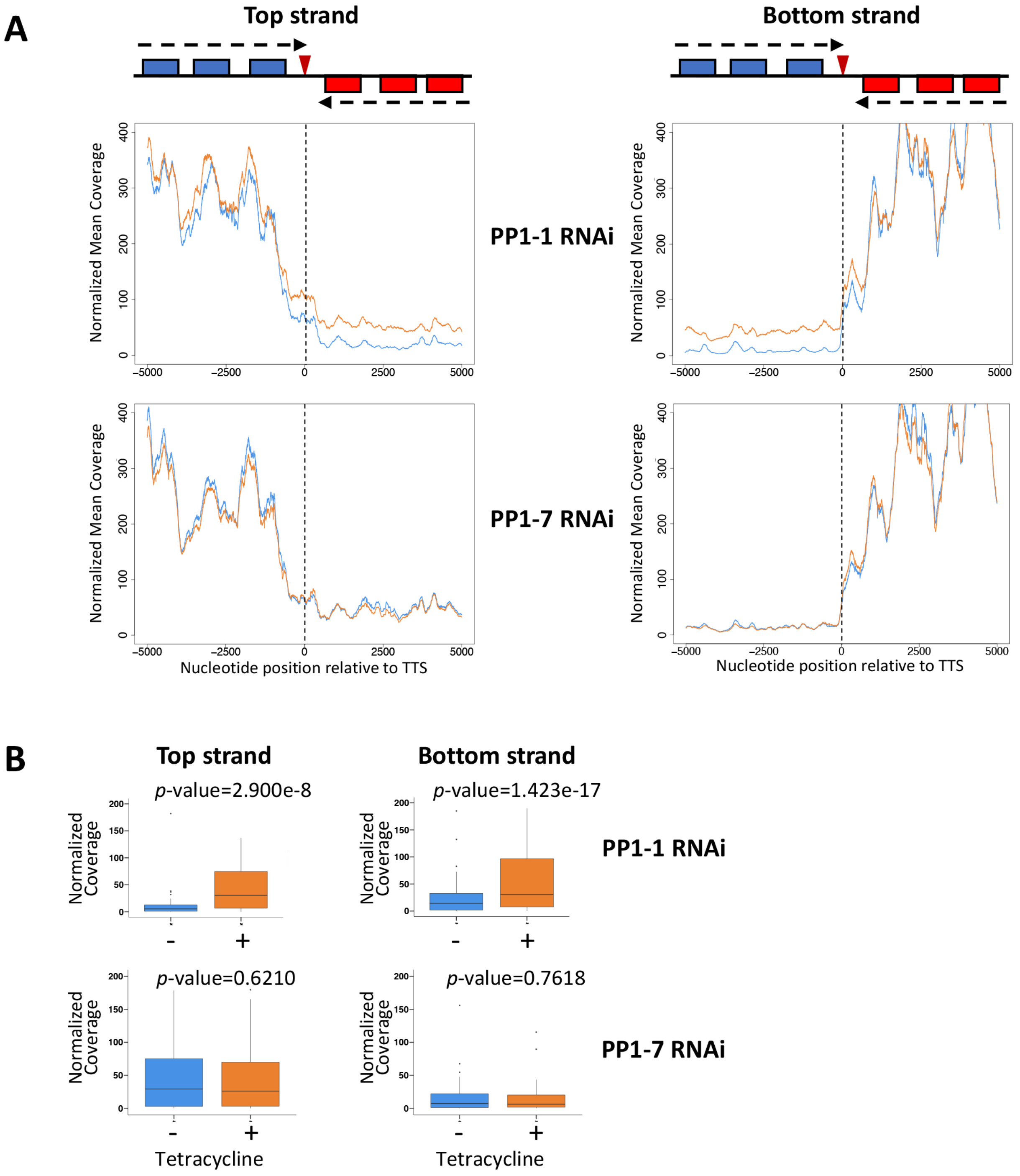
A
Top strand
Bottom strand
PP1-1 RNAi
PP1-7 RNAi
Normalized Mean Coverage
Nucleotide position relative to TTS
B
Top strand
Bottom strand
p-value=2.900e-8
p-value=1.423e-17
p-value=0.6210
p-value=0.7618
Normalized Coverage
PP1-1 RNAi
PP1-7 RNAi
Tetracycline

Figure S6

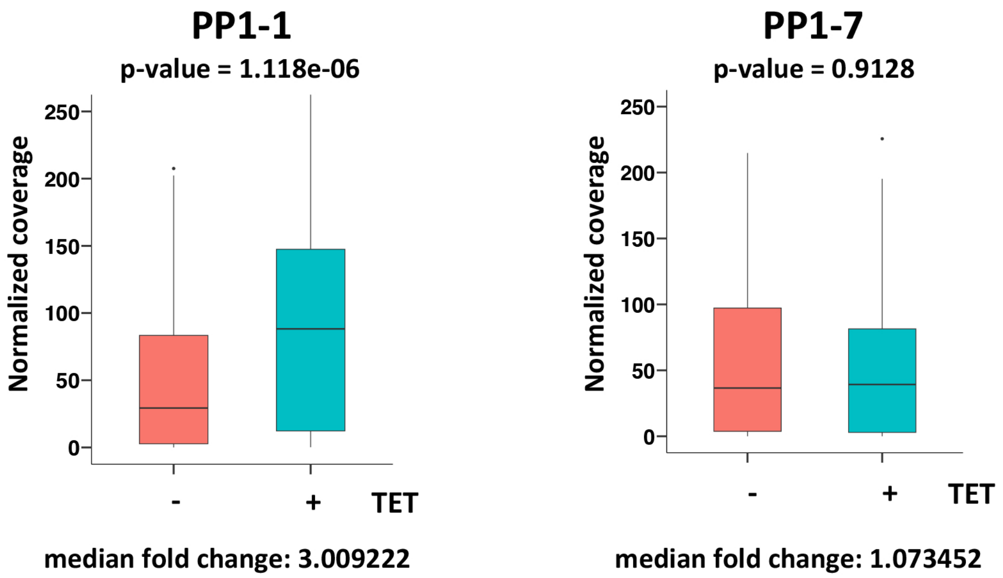

PP1-1
p-value = 1.118e-06
Normalized coverage
250
200
150
100
50
0
-
+
TET
median fold change: 3.009222
PP1-7
p-value = 0.9128
Normalized coverage
250
200
150
100
50
0
-
+
TET
median fold change: 1.073452


Figure S7

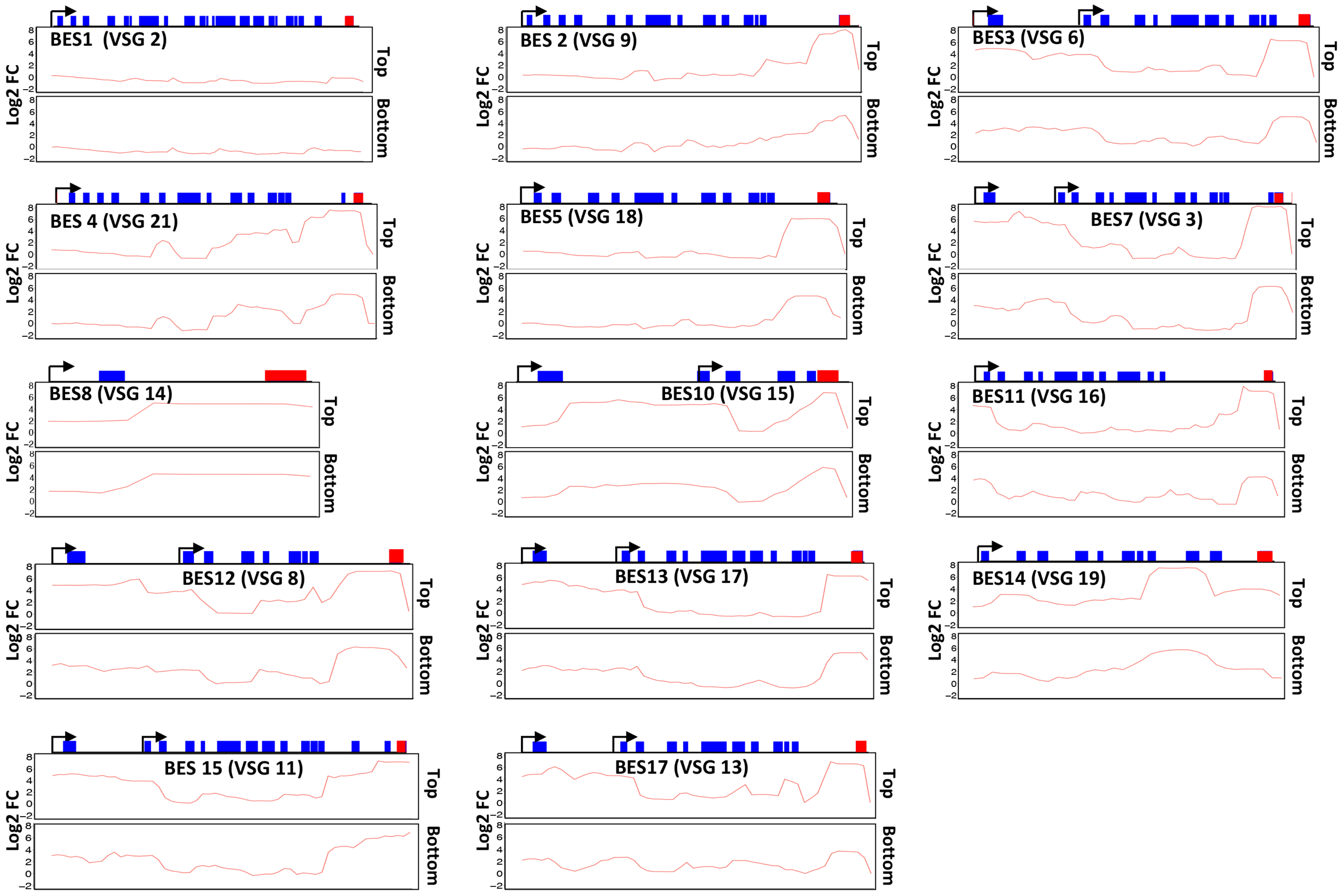
BES1 (VSG 2)
BES 2 (VSG 9)
BES3 (VSG 6)
BES 4 (VSG 21)
BES5 (VSG 18)
BES7 (VSG 3)
BES8 (VSG 14)
BES10 (VSG 15)
BES11 (VSG 16)
BES12 (VSG 8)
BES13 (VSG 17)
BES14 (VSG 19)
BES 15 (VSG 11)
BES17 (VSG 13)
Log2 FC
Top
Bottom
8
6
4
2
0
−2

**Figure S8**

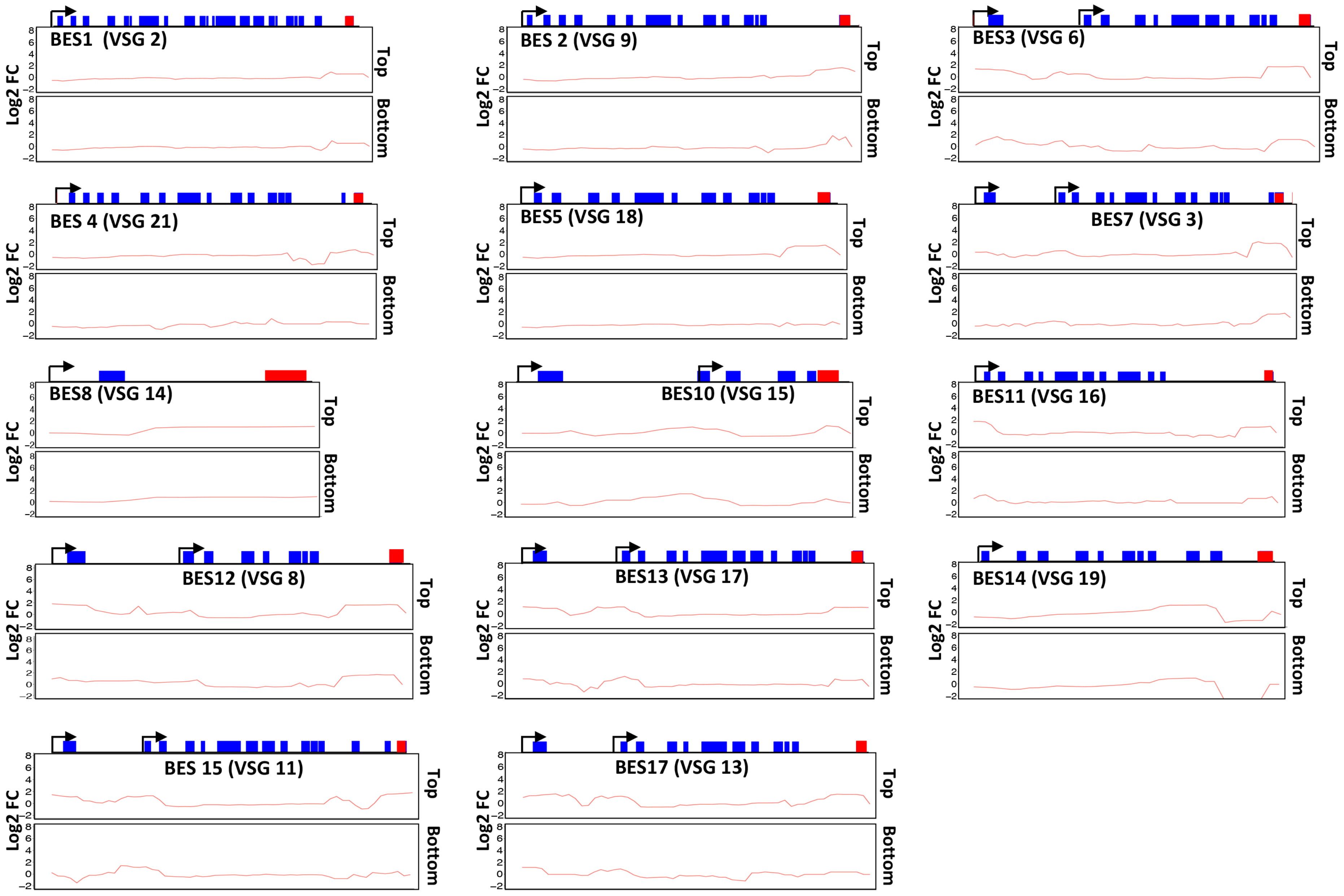
BES1 (VSG 2)
BES 2 (VSG 9)
BES3 (VSG 6)
BES 4 (VSG 21)
BES5 (VSG 18)
BES7 (VSG 3)
BES8 (VSG 14)
BES10 (VSG 15)
BES11 (VSG 16)
BES12 (VSG 8)
BES13 (VSG 17)
BES14 (VSG 19)
BES 15 (VSG 11)
BES17 (VSG 13)
Log2 FC
Top
Bottom
8
6
4
2
0
−2

**Figure S9**

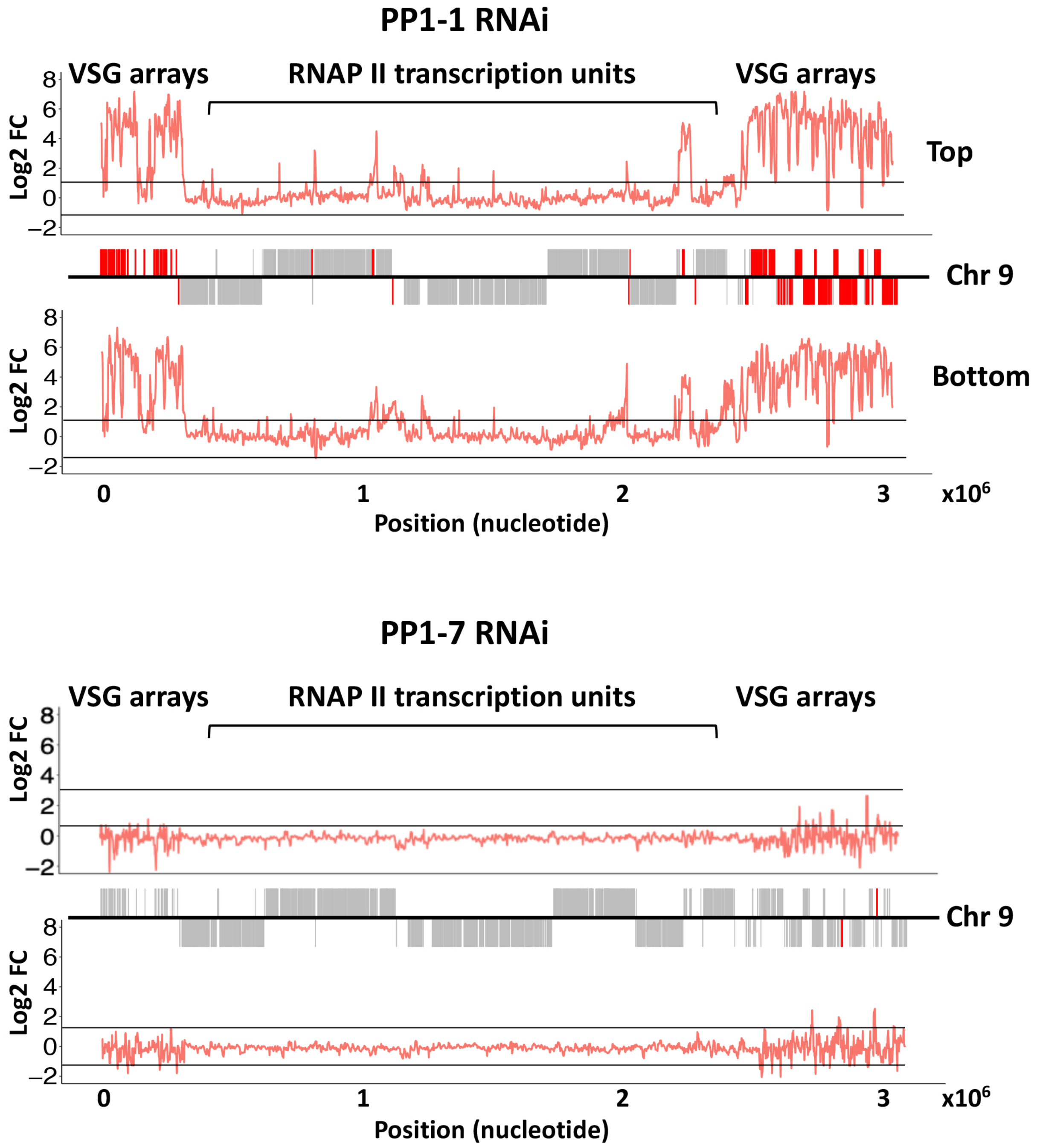
PP1-1 RNAi
VSG arrays
RNAP II transcription units
VSG arrays
Log2 FC
Top
Chr 9
Bottom
Position (nucleotide)
PP1-7 RNAi
VSG arrays
RNAP II transcription units
VSG arrays
Log2 FC
Chr 9
Position (nucleotide)

Figure S10

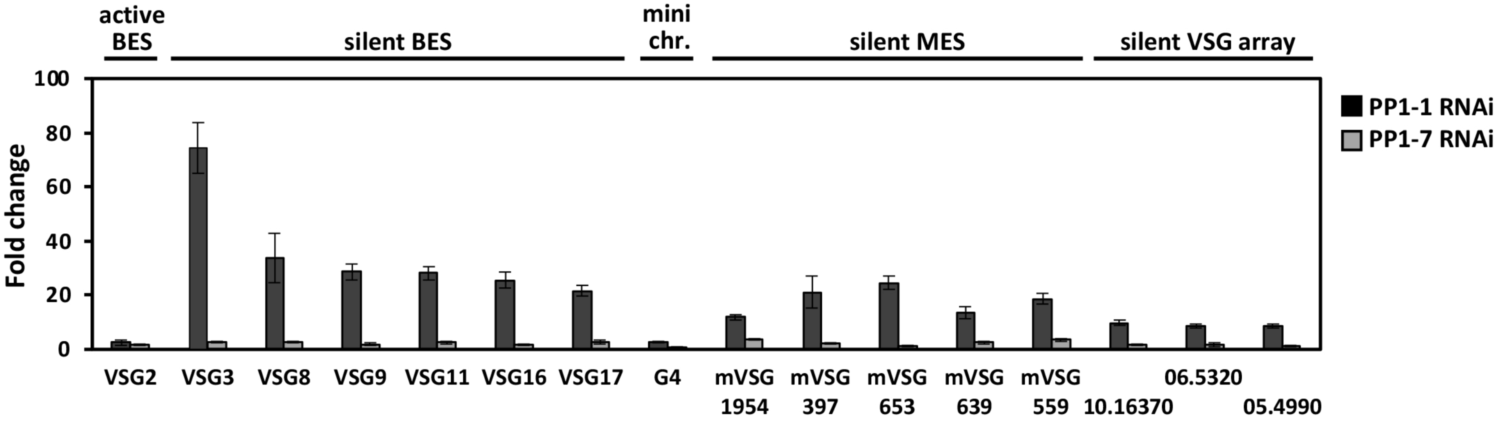

active BES
silent BES
mini chr.
silent MES
silent VSG array
PP1-1 RNAi
PP1-7 RNAi
Fold change
100
80
60
40
20
0
VSG2
VSG3
VSG8
VSG9
VSG11
VSG16
VSG17
G4
mVSG 1954
mVSG 397
mVSG 653
mVSG 639
mVSG 559
10.16370
06.5320
05.4990


**Figure S11**

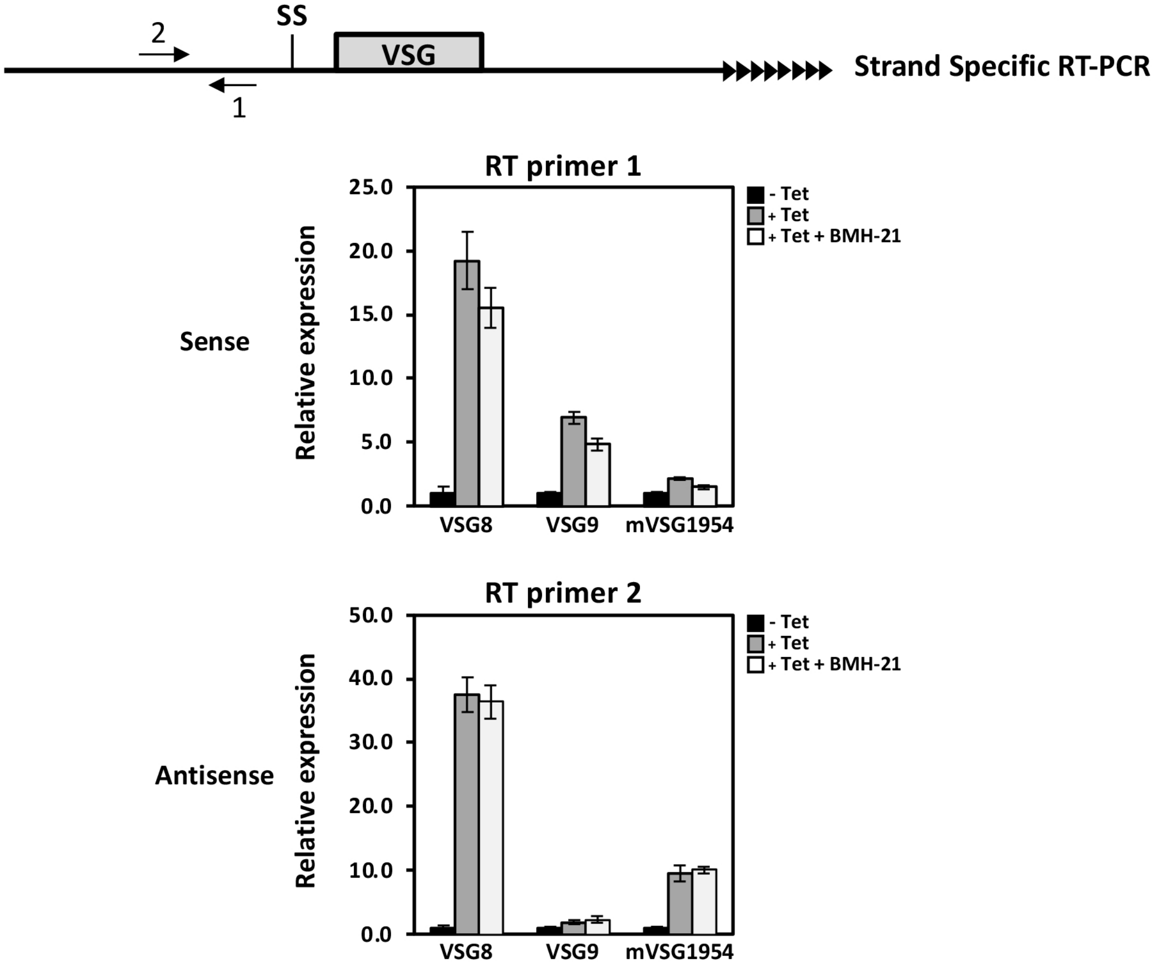
2
SS
VSG
1
Strand Specific RT-PCR
Sense
Antisense
RT primer 1
RT primer 2
Relative expression
25.0
20.0
15.0
10.0
5.0
0.0
50.0
40.0
30.0
20.0
10.0
VSG8
VSG9
mVSG1954
- Tet
+ Tet
+ Tet + BMH-21

Figure S12

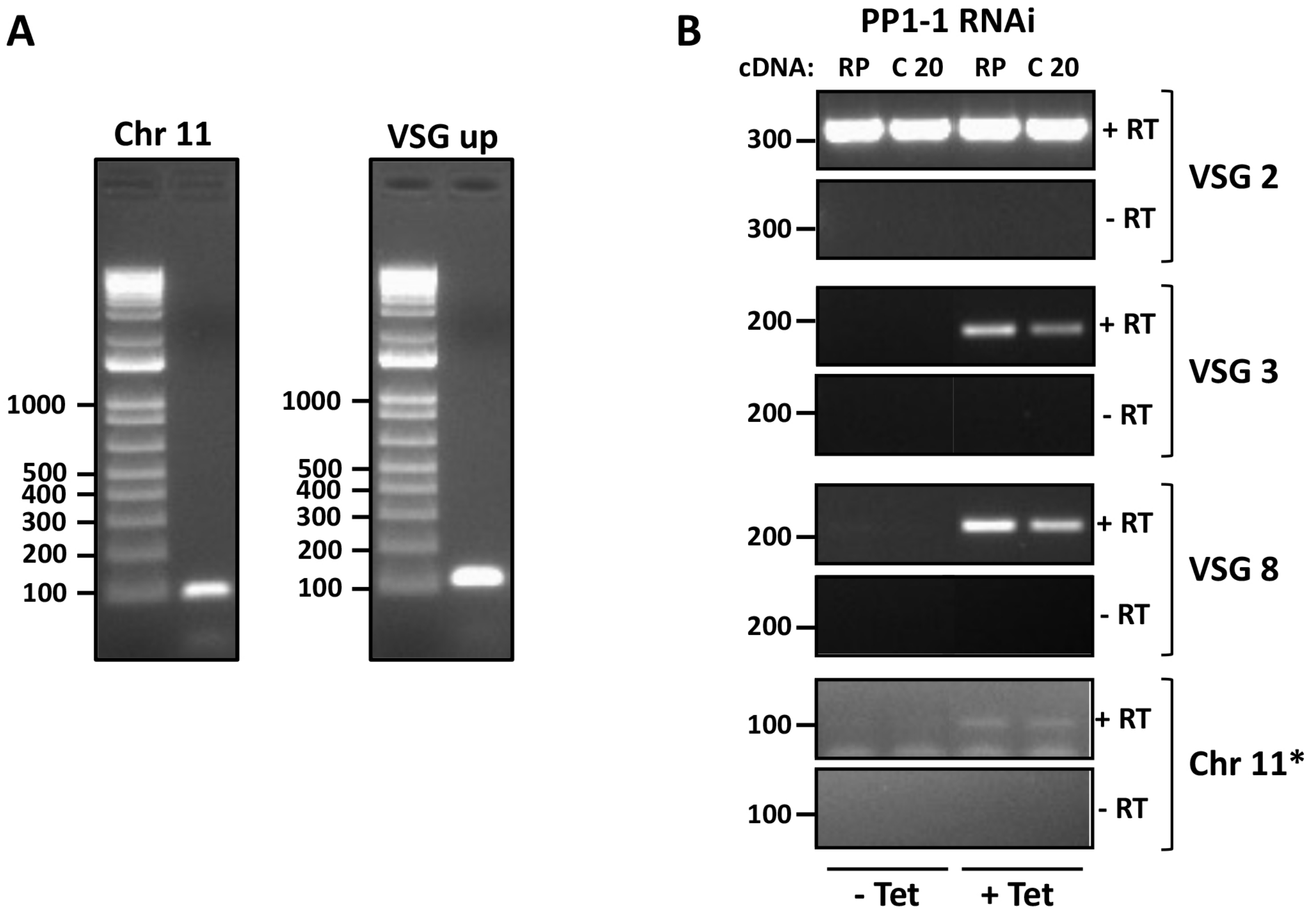
A
Chr 11
VSG up
1000
500
400
300
200
100
B
PP1-1 RNAi
cDNA: RP C 20 RP C 20
+ RT
- RT
VSG 2
VSG 3
VSG 8
Chr 11*
- Tet
+ Tet

Figure S13

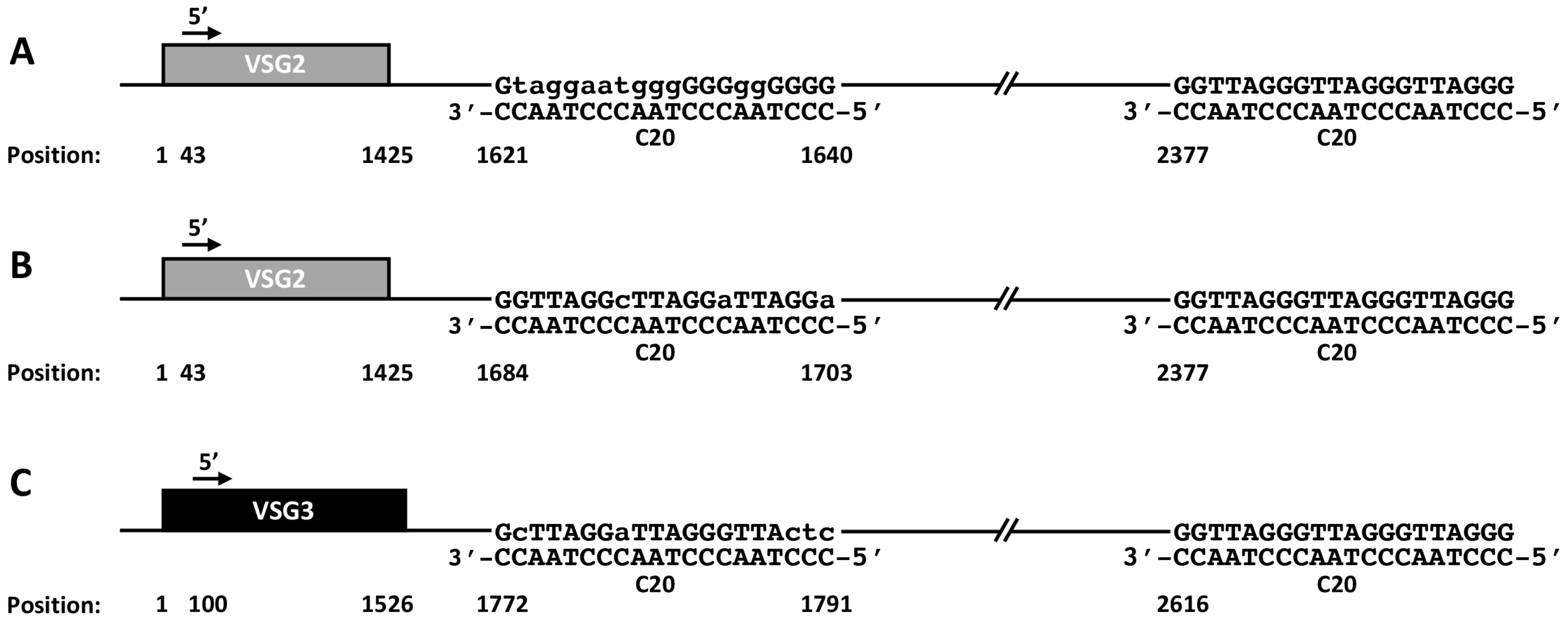
A
5′
VSG2
Gtaggaatgggg GGGggGGGG
3′-CCAATCCCAATCCCAATCCC-5′
C20
GGTTAGGGTTAGGGTTAGGG
3′-CCAATCCCAATCCCAATCCC-5′
C20
Position: 1 43 1425 1621 1640 2377
B
5′
VSG2
GGTTAGGcTTAGGaTTAGGa
3′-CCAATCCCAATCCCAATCCC-5′
C20
GGTTAGGGTTAGGGTTAGGG
3′-CCAATCCCAATCCCAATCCC-5′
C20
Position: 1 43 1425 1684 1703 2377
C
5′
VSG3
GcTTAGGaTTAGGGTTActc
3′-CCAATCCCAATCCCAATCCC-5′
C20
GGTTAGGGTTAGGGTTAGGG
3′-CCAATCCCAATCCCAATCCC-5′
C20
Position: 1 100 1526 1772 1791 2616

**Figure S14**

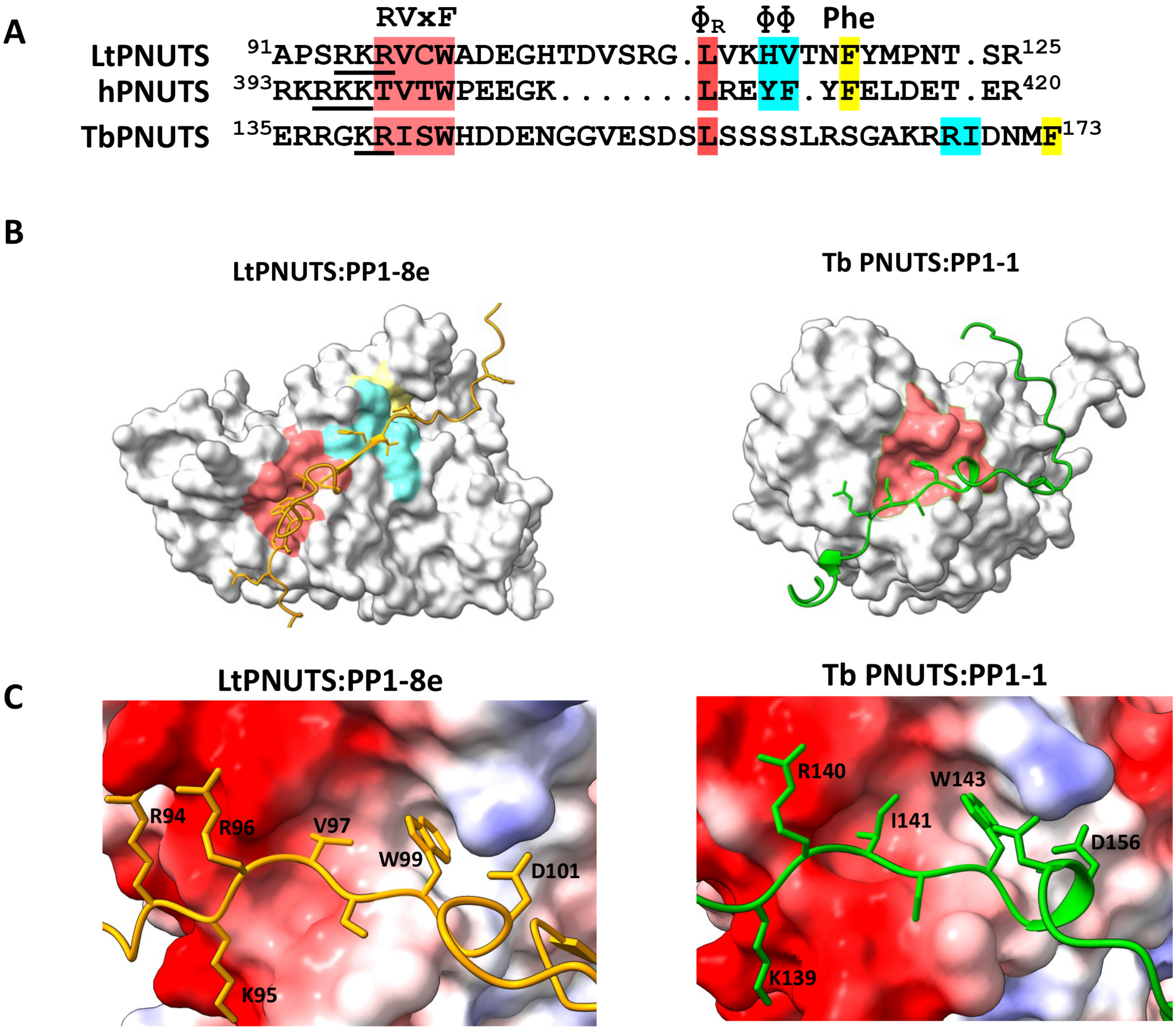

A
RVxF
ΦR
ΦΦ
Phe
LtPNUTS 91APSRKRVCWADEGHTDVSRG.LVKHVTNFYMPNT.SR125
hPNUTS 393RKRKKTVTWPEEGK........LREYF.YFELDET.ER420
TbPNUTS 135ERRGKRISWHDDENGGVESDSLSSSSLRSGAKRRIDNMF173
B
LtPNUTS:PP1-8e
Tb PNUTS:PP1-1
C
LtPNUTS:PP1-8e
R94
R96
V97
W99
D101
K95
Tb PNUTS:PP1-1
R140
W143
I141
D156
K139


Figure S15

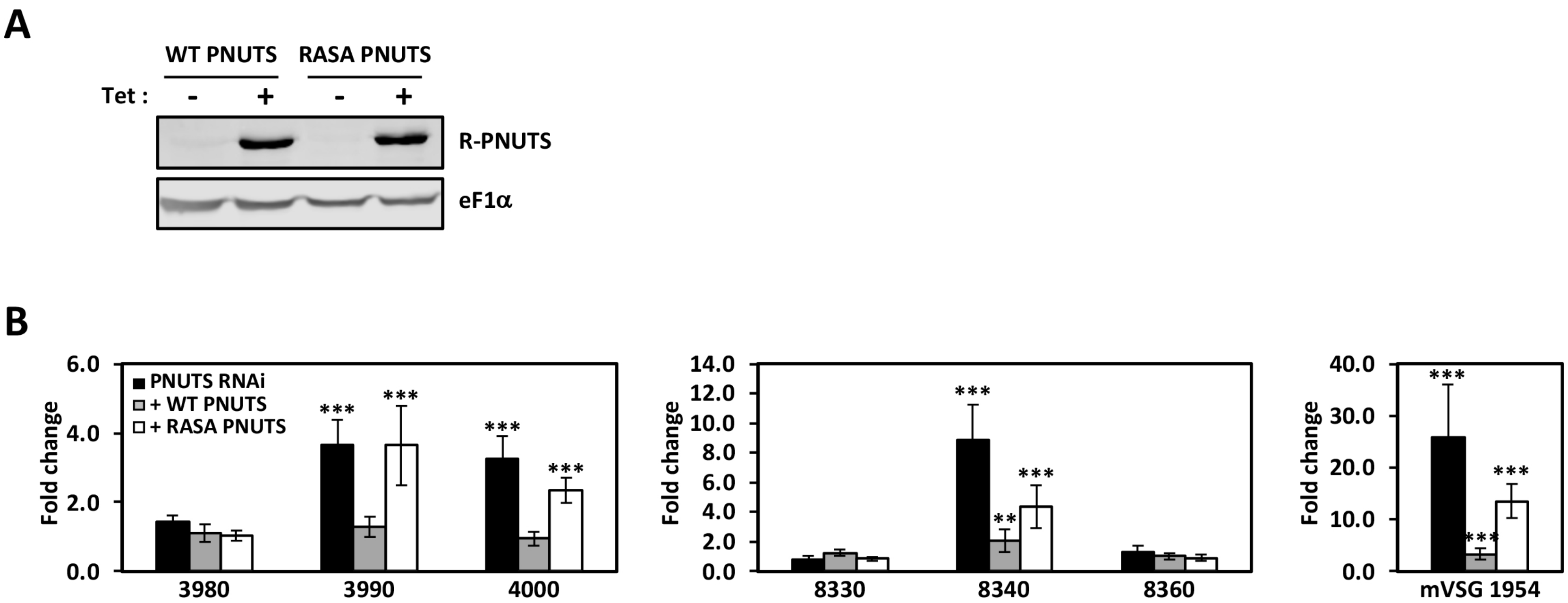
A
WT PNUTS
RASA PNUTS
Tet :
-
+
-
+
R-PNUTS
eF1α
B
6.0
4.0
2.0
0.0
Fold change
PNUTS RNAi
+ WT PNUTS
+ RASA PNUTS
***
3980
3990
4000
14.0
12.0
10.0
8.0
6.0
4.0
2.0
0.0
**
8330
8340
8360
40.0
30.0
20.0
10.0
0.0
mVSG 1954

Figure S16
